# Dose-dependent modeling of combinatorial drug responses stratifies patient survival and reveals therapeutic vulnerabilities in precision oncology

**DOI:** 10.64898/2026.04.16.718332

**Authors:** Kohei Ota, Takumi Ito, Hideyuki Shimizu

## Abstract

A substantial proportion of cancer patients fail to benefit from their prescribed combination regimens, yet identifying superior alternatives from the vast pharmacological space prior to treatment failure remains an unsolved clinical challenge. Existing computational approaches either rely on multi-omics profiles unavailable in standard oncological practice or reduce drug efficacy to scalar metrics that discard the dose-dependent resolution essential for therapeutic optimization. Here, we present XACT, a hierarchical deep learning framework that reconstructs full dose-dependent drug responses for both monotherapy and drug combinations using only clinically accessible transcriptomic profiles. By leveraging an asymmetric X-Linear Attention mechanism that models second-order interactions between molecular drug substructures and intracellular signaling pathway activities, XACT captures concentration-dependent pharmacodynamics with state-of-the-art accuracy and generalizability to unseen transcriptomic landscapes. When applied to the TCGA pan-cancer cohort, XACT-derived resistance scores were significantly associated with clinical treatment outcomes and stratified overall survival as the strongest independent prognostic factor after multivariate adjustment for tumor stage and cancer type. Systematic virtual screening revealed therapeutic vulnerabilities and nominated alternative regimens for treatment-refractory sarcoma and pancreatic adenocarcinoma. These results establish XACT as a scalable, interpretable, and clinically translatable framework that advances precision oncology from computational prediction toward data-driven therapeutic prescription.

## Introduction

Cancer remains the leading cause of mortality worldwide, and despite decades of therapeutic development, a substantial proportion of patients fail to achieve durable responses to their prescribed treatment regimens^1,2^. The clinical benefit conferred by any given regimen varies profoundly across individuals, driven by the molecular heterogeneity of tumors and the clonal evolutionary dynamics that underlie acquired drug resistance^3–6^. For a given patient whose tumor does not respond to the standard regimen, a superior alternative may exist within the approved pharmacological space, and the central challenge is to identify it before treatment failure occurs.

Combination chemotherapy, which co-deploys agents with distinct mechanisms of action to induce synergistic cytotoxicity and suppress resistant clones, represents the backbone of modern oncological practice^7^. However, the combinatorial search space of approved drugs is vast, and experimentally screening all possible pairs and concentration ratios for each patient is prohibitively expensive and time-consuming for routine clinical application. This systematic bottleneck has driven the development of computational approaches aimed at predicting drug responses *in silico*, ideally leveraging large-scale pharmacogenomic databases such as the Genomics of Drug Sensitivity in Cancer (GDSC)^8^ to guide therapeutic decision-making and accelerate the discovery of superior regimens for refractory patients.

Despite substantial progress, current computational efforts^9^ are constrained by three fundamental limitations that hinder their clinical translation. First, while a substantial body of monotherapy response prediction models exists, the overwhelming majority predict only a scalar IC₅₀ value per drug-cell pair and are thus unable to reconstruct the continuous dose-dependent viability trajectory across the concentration range^10–13^. A small subset of models extend this capability to full dose-response curve prediction^14^, yet even these remain confined to monotherapy contexts. Models capable of predicting combinatorial drug responses are considerably rarer, and this gap is clinically significant given that the overwhelming majority of oncological regimens involve two or more agents. Furthermore, virtually all reduce combinatorial efficacy to a scalar synergy score rather than a dose-dependent viability surface^15–17^, discarding the pharmacodynamic resolution necessary for rational dose optimization. No existing model is capable of predicting the holistic response to regimens comprising three or more agents, despite the fact that many standard oncological regimens involve combinations of this complexity. Rational clinical decision-making requires knowledge of the full pharmacodynamic trajectory across the relevant concentration space, not a single summary statistic. Second, state-of-the-art models frequently require extensive multi-omics profiles encompassing somatic mutations, copy number variations, and DNA methylation^10,18,19^, data modalities that remain unavailable for the majority of patients in routine oncological practice. Static genomic markers are furthermore inherently unable to capture the dynamic functional remodeling of cellular signaling pathways that underlies drug resistance acquisition. Transcriptomic signatures derived from bulk RNA sequencing represent a more accessible and functionally descriptive alternative, providing a real-time readout of cellular state and pathway activity while circumventing the operational bottlenecks of multi-omics integration. Third, and most critically, existing models are primarily optimized for preclinical cell-line datasets and have rarely been validated against patient outcome data, a failure attributable to biological noise and intratumoral heterogeneity. As a consequence, even the most accurate preclinical models have yet to demonstrate utility in stratifying patient outcomes or guiding treatment selection in real-world oncological practice.

To bridge this translational gap, we present XACT (X-Linear Attention for Combination Therapy), a hierarchical deep learning framework comprising three coupled modules. XACT-Single first reconstructs full dose-dependent monotherapy responses from clinically accessible transcriptomic profiles alone, establishing robust pharmacological representations that serve as the foundation for combinatorial modeling. XACT-Dual then extends this architecture into a weight-shared Siamese framework augmented with asymmetric cross-branch attention to predict dose-dependent combinatorial drug responses across the full two-dimensional concentration space, capturing the directional pharmacological modulation between co-administered agents. XACT-Clinical subsequently adapts the combinatorial framework to the clinical domain, generating resistance scores that stratify patient outcomes in real-world cohorts. Prospective experimental validation confirmed that XACT predictions generalize to cancer types and mechanistic contexts entirely absent from training data. Systematic virtual screening revealed therapeutic vulnerabilities and nominated alternative regimens for treatment-refractory sarcoma and pancreatic adenocarcinoma. Crucially, XACT-derived resistance scores emerged as the strongest independent prognostic factor in the TCGA pan-cancer cohort after multivariate adjustment for tumor stage and cancer type. Together, these findings position XACT as a clinically actionable framework that bridges the divide between pharmacogenomic discovery and patient-level therapeutic decision-making.

## Results

### XACT-Single predicts dose-dependent drug response with high accuracy and generalizability across unseen transcriptomic landscapes

XACT comprises three hierarchically coupled modules designed to bridge the translational gap between preclinical pharmacogenomics and clinical practice. The foundational module, XACT-Single, is grounded in the biological principle that accurate monotherapy response prediction provides the pharmacological substrate necessary for modeling combinatorial effects (**Fig. 1A**). Drug representations integrate complementary structural and functional perspectives, with topological features extracted from the molecular graph via a Graph Isomorphism Network (GIN)^20^ to capture chemical architecture, while functional bioactivity signatures derived from the Chemical Checker^21^ framework encode known pharmacological profiles across multiple levels of biological organization. Cellular context is represented as pathway-level activity scores computed by ssGSEA^22^ on bulk RNA-seq data, encoding the tumor transcriptome as a landscape of dynamic signaling pathway activations rather than a static vector of gene expression counts. These heterogeneous modalities are integrated via an X-Linear Attention mechanism^23^ that explicitly models second-order interactions between molecular drug substructures and intracellular signaling pathways.

**Fig. 1.**
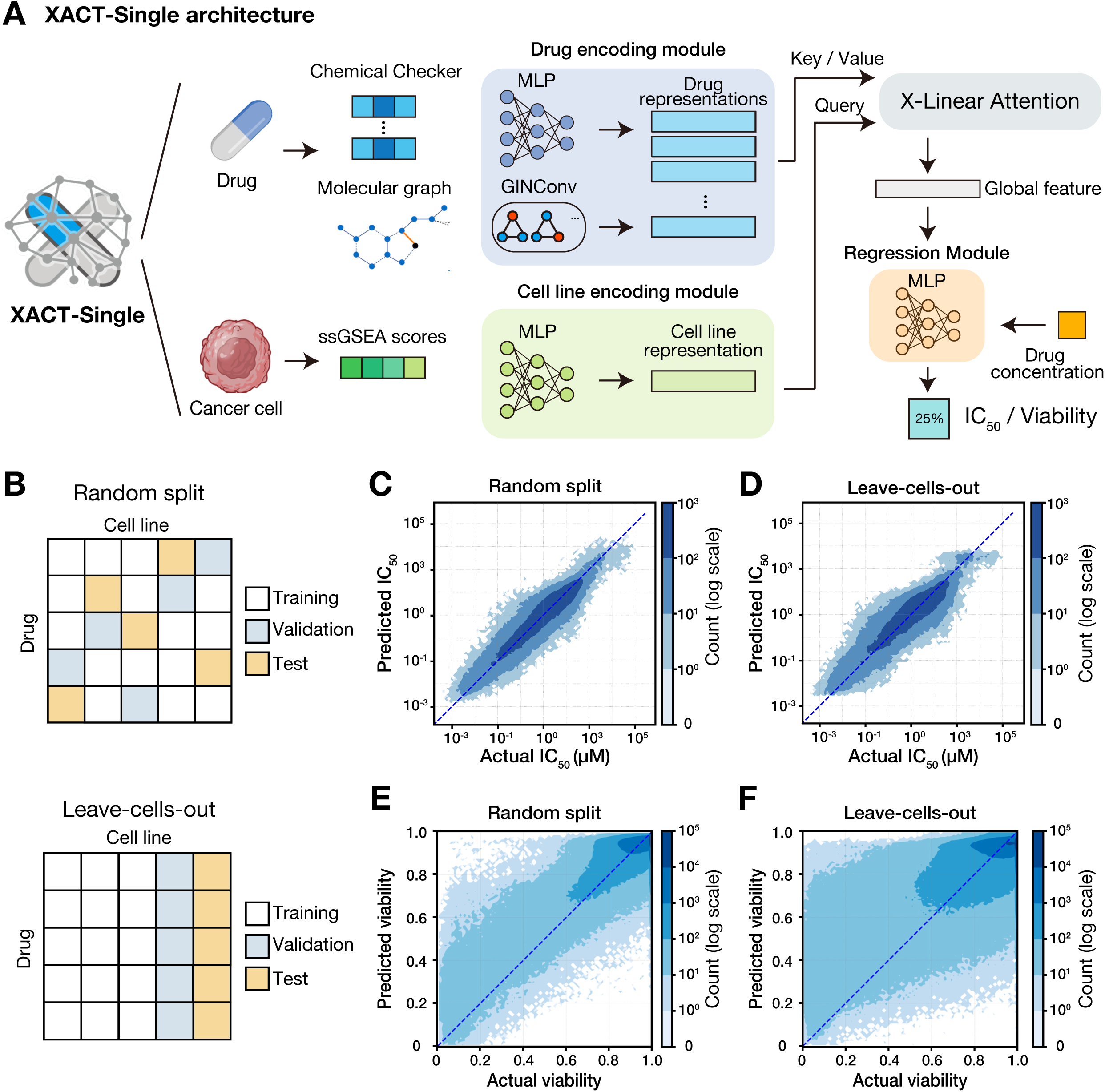
Architecture and validation strategy of XACT-Single. **(A)** Architecture of the XACT-Single monotherapy response prediction module. Each drug is encoded through two parallel branches capturing complementary aspects of molecular identity: topological features are extracted from the molecular graph via a Graph Isomorphism Network (GIN), while functional bioactivity signatures are derived from the Chemical Checker framework. Cellular context is encoded as pathway-level activity scores computed by ssGSEA on bulk RNA-seq data and projected into the shared latent space through a dedicated MLP. These heterogeneous modalities are integrated through an X-Linear Attention mechanism, in which the cellular embedding serves as a dynamic query that attends over the multi-view drug representation to produce a global drug-cell interaction feature. Drug concentration is introduced via late fusion, concatenated with the global feature vector immediately prior to the final regression layer, ensuring that the upstream feature extraction remains concentration-independent. **(B)** Validation strategy. Two cross-validation schemes of increasing stringency are employed: random split, which assesses predictive accuracy on held-out drug-cell pairs drawn from the same distribution as the training data, and leave-cells-out, which withholds entire cell lines from training to evaluate generalization to unseen transcriptomic landscapes, directly mimicking the clinical scenario of predicting drug response in a previously uncharacterized patient. **(C–D)** Two-dimensional density plots illustrating the concordance between XACT-Single predictions and experimentally measured IC₅₀ values under random split **(C)** and leave-cells-out **(D)** conditions. Data points are aggregated into 100×100 two-dimensional bins, with color encoding the local density of observations on a logarithmic scale such that darker regions indicate higher concentrations of predictions and lighter regions represent sparse outliers. The dashed diagonal line represents perfect concordance. Both axes are plotted on a logarithmic scale in micromolar units. **(E–F)** Two-dimensional density plots illustrating concordance between predicted and observed fractional cell viability under random split **(E)** and leave-cells-out **(F)** conditions. The tight clustering of predictions along the diagonal identity line across all four panels demonstrates that XACT-Single maintains high predictive fidelity under both interpolation (random split) and extrapolation (leave-cells-out) conditions.

To establish the translational validity of XACT-Single, a hierarchical validation strategy of increasing stringency was implemented (**Fig. 1B**). Random split cross-validation assessed predictive accuracy under interpolation conditions, whereas the more stringent leave-cells-out scheme withheld entire cell lines from training to evaluate generalization to transcriptomic landscapes entirely absent from the training distribution, directly mimicking the clinical scenario of predicting drug response in a previously uncharacterized patient^24^.

Evaluated on the GDSC dataset comprising 118,896 IC₅₀ measurements across 959 cancer cell lines and 162 compounds, XACT-Single demonstrated exceptional predictive accuracy, achieving a Pearson correlation coefficient (PCC) of 0.923 ± 0.007 under random split (**Fig. 1C**). Crucially, the model maintained high generalizability under the more demanding leave-cells-out task (PCC = 0.895 ± 0.002), with predictions distributed tightly along the identity line across the full dynamic range of the sensitivity spectrum (**Fig. 1D**). This less than 3% degradation in PCC across validation schemes indicates that XACT-Single learns transferable pharmacological rules rather than memorizing cell-line-specific statistical patterns. Benchmarking against three established computational methods (CDS^10^, PathDNN^11^, and HiDRA^13^) together with a baseline multilayer perceptron (MLP) model confirmed that XACT-Single outperformed all competitors across every evaluated metric under the rigorous leave-cells-out condition (*t*-test, *p* < 0.05; **Supplementary Tables 1 and 2**). Beyond quantitative performance, mechanistic interrogation via SHAP^25^ and GNNExplainer^26^ confirmed that these predictions are driven by pharmacologically coherent features. Specifically, KRAS and RAF signaling emerged as dominant transcriptomic determinants of sensitivity to the MEK inhibitor trametinib, consistent with canonical MEK pathway dependencies^27,28^. Furthermore, the allosteric pharmacophore of trametinib prioritized as the structural basis of predicted potency^29^ (**Supplementary Fig. 1**), demonstrating that XACT-Single has internalized biologically valid structure-activity and genotype-sensitivity relationships rather than exploiting dataset-specific statistical artifacts.

Whereas existing monotherapy models predominantly predict a single IC₅₀ scalar per drug-cell pair and thus provide no information on how cellular response evolves across the concentration range^10,11,13,19^, XACT-Single is explicitly architected to reconstruct full dose-dependent cell viability curves, a capability essential for delineating therapeutic windows and characterizing the continuous pharmacodynamic relationship between drug concentration and cellular response. Predicted viability trajectories closely matched experimental measurements under both random split (**Fig. 1E**) and leave-cells-out (**Fig. 1F**) conditions, with consistently low root-mean-square error (RMSE) across all evaluated drug-cell line pairs (**Supplementary Table 3**).

### XACT-Single accurately reconstructs dose-response profiles across diverse cancer types and mechanistic contexts

To illustrate the fidelity of these reconstructions across pharmacodynamically distinct contexts, we examined three representative drug-cell line pairs: paclitaxel in MDA-MB-231 breast cancer cells (**Fig. 2A**), cisplatin in A549 non-small cell lung cancer cells (**Fig. 2B**), and gefitinib in the same A549 cell line (**Fig. 2C**). Predictions closely recapitulated the experimentally observed pharmacodynamic trajectories, including both the concentration range over which viability declined and the asymptotic survival fraction at saturating doses. A549 cells harbor a *KRAS* mutation that confers intrinsic resistance to EGFR inhibition^30^, and gefitinib treatment accordingly produced minimal reduction in cell viability across the tested concentration range, representing a resistant pharmacodynamic profile that XACT-Single correctly anticipated. This capacity for accurate dose-response reconstruction was consistent across a broad panel of pharmacologically distinct drug classes and cancer cell lineages, including breast, lung, kidney, melanoma, colorectal, and hematological malignancies, as demonstrated by nine representative examples (**Supplementary Fig. 2**).

**Fig. 2.**
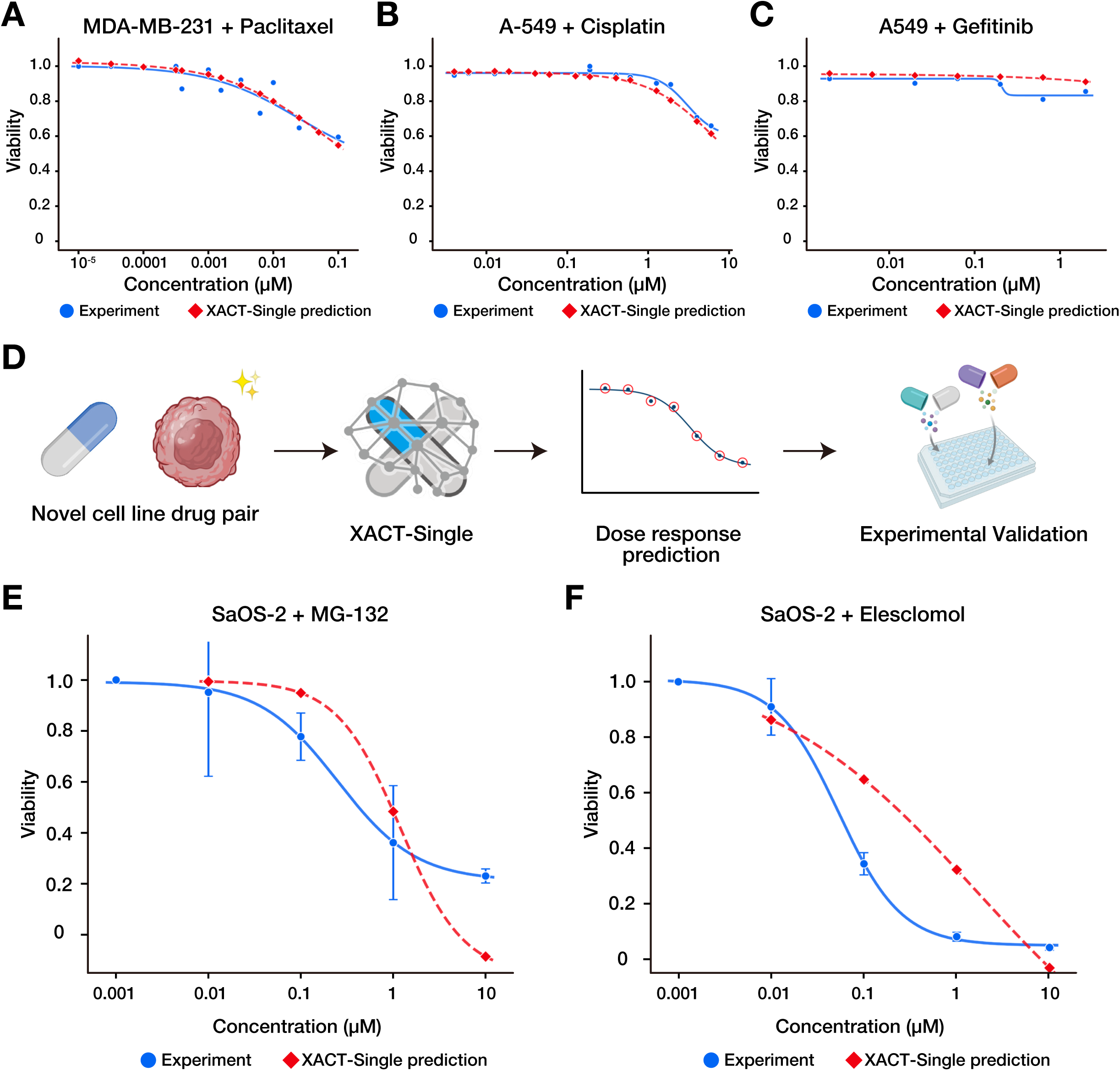
Experimental validation of XACT-Single predictions in novel cell line and drug contexts. **(A-C)** Predicted and experimentally observed dose-response curves for three representative drug-cell line pairs drawn from the GDSC dataset, spanning distinct cancer types and pharmacological response profiles: MDA-MB-231 treated with paclitaxel (Breast, TNBC) **(A)**, A549 treated with cisplatin (Lung, NSCLC) **(B)**, and A549 treated with gefitinib (Lung, NSCLC) **(C)**. In each panel, the horizontal axis represents drug concentration in micromolar units on a logarithmic scale and the vertical axis represents fractional cell viability. Experimentally measured values are shown as blue circles and XACT-Single predictions as red diamonds, each fitted with a four-parameter Hill equation. **(D)** For any user-defined drug-cell line combination, the model generates a predicted dose-response curve that can be directly subjected to experimental validation, enabling hypothesis-driven prioritization of novel therapeutic contexts prior to laboratory testing. **(E–F)** Prospective experimental validation of XACT-Single predictions for two drug-cell line pairs entirely absent from the training data: the proteasome inhibitor MG-132 in the osteosarcoma cell line SaOS-2 **(E)** and the copper-chelating agent elesclomol in SaOS-2 **(F)**. XACT-Single predictions are shown as a red dashed line with diamond markers alongside experimentally measured cell viability values shown as blue solid lines with circles, representing the mean ± standard deviation across three biological replicates. The close agreement between predicted and observed dose-response profiles across both compounds demonstrates the prospective generalization capability of XACT-Single to cancer types and mechanistic contexts not encountered during training.

Osteosarcoma represents a malignancy of profound unmet need^31^, characterized by limited second-line therapeutic options and poor prognosis in metastatic disease. To demonstrate that XACT-Single can prospectively identify pharmacologically active compounds for such refractory cancers without prior experimental data, we selected SaOS-2, an osteosarcoma cell line entirely absent from the GDSC training dataset, as a challenging test case (**Fig. 2D**). Predictions were generated for two mechanistically distinct compounds: MG-132, a proteasome inhibitor^32^, and elesclomol, a copper-chelating agent that induces cuproptosis^33^. Independent cell viability assays confirmed close agreement between predicted and experimentally measured dose-response profiles across the tested concentration range for both compounds (**Fig. 2E, F**), providing prospective experimental evidence that the pharmacological representations learned from large-scale cell line data encode transferable drug sensitivity rules that generalize to cancer types and mechanistic contexts not encountered during training.

### State-of-the-art precision in combinatorial response prediction with XACT-Dual

Building upon the pharmacological representations established during monotherapy training, XACT-Dual was developed to predict dose-dependent viability for drug pairs by extending the XACT-Single architecture into a weight-shared Siamese framework^34^ augmented with asymmetric cross-branch attention (**Fig. 3A**). The monotherapy encoders were held fixed as stable feature extractors, preserving the fundamental pharmacological representations of each drug and preventing overfitting to the comparatively sparse combinatorial data. The core innovation is an asymmetric cross-branch attention mechanism in which the global representation of one drug serves as a query that interrogates the multi-view feature representation of its partner as key and value^23^. This design explicitly models the directional modulation of one drug’s mechanism of action by the other, mathematically formalizing the context-dependency of pharmacological synergy that symmetric architectures are inherently unable to represent.

**Fig. 3.**
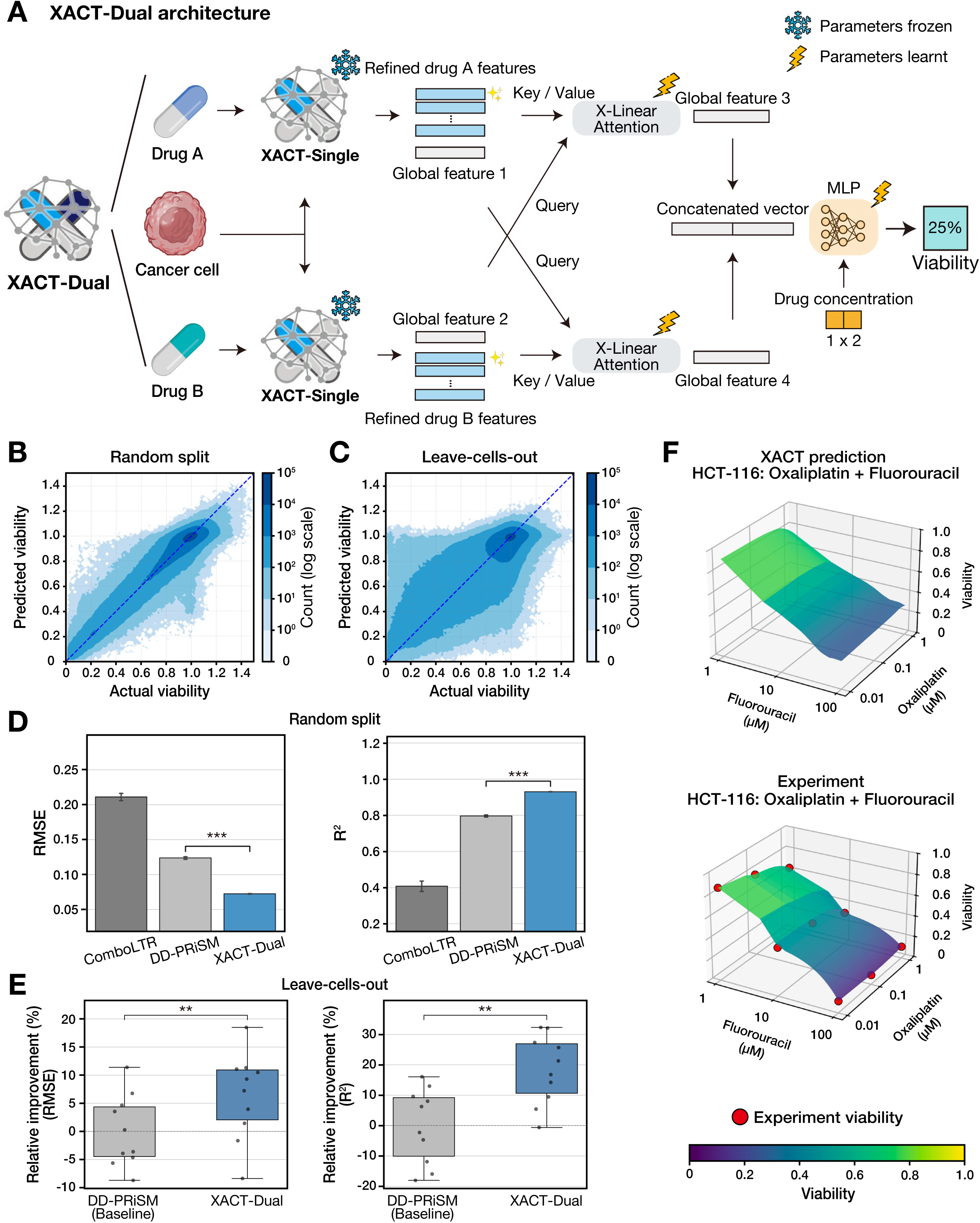
Architecture, benchmarking, and dose-response surface reconstruction of XACT-Dual. **(A)** Architecture of the XACT-Dual combinatorial response prediction module. The XACT-Single encoder is instantiated as a weight-shared Siamese network framework, with all monotherapy parameters held fixed to preserve the pharmacological representations established during single-drug training. Each branch independently processes one drug alongside the shared cell line representation to produce a global interaction vector and a multi-view attention map. Drug-drug interactions are modeled by two trainable cross-branch X-Linear Attention blocks, in which the global feature of each drug serves as a query that attends over the multi-view representation of its combination partner, explicitly encoding the directional pharmacological modulation between co-administered agents. The resulting cross-attended features are concatenated with the concentrations of both drugs and passed through a final MLP to predict dose-dependent combinatorial cell viability. **(B–C)** Two-dimensional density plots illustrating the concordance between XACT-Dual predictions and experimentally measured two-drug cell viability under random split **(B)** and leave-cells-out **(C)** conditions. Data points are aggregated into 100×100 two-dimensional bins, with color encoding the local density of observations on a logarithmic scale such that darker regions indicate higher concentrations of predictions and lighter regions represent sparse outliers. The dashed diagonal line represents perfect concordance. **(D)** Predictive performance of XACT-Dual, DD-PRiSM, and ComboLTR under random split cross-validation, quantified by Root Mean Square Error (RMSE) and Coefficient of Determination (R²). Bar height represents the mean across cross-validation folds and error bars indicate standard deviation. Statistical significance of pairwise differences was assessed by two-sided t-test (\*\*\**p* < 0.001). **(E)** Relative improvement of XACT-Dual over DD-PRiSM under leave-cells-out cross-validation, quantified as the percentage of remaining room for improvement, defined as the gap between DD-PRiSM performance and the theoretical optimum. Each data point represents one cross-validation fold and the box represents the interquartile range. Statistical significance was assessed by two-sided *t*-test (\*\**p* < 0.01). **(F)** Representative three-dimensional dose-response surface for the combination of oxaliplatin and fluorouracil in the HCT-116 colorectal cancer cell line, showing the XACT-Dual prediction (top) and the experimentally determined surface (bottom). Red circles indicate measured viability values at specific drug concentration pairs. Surface color encodes fractional cell viability on a continuous scale from 0 (dark purple, complete cell death) to 1.0 (yellow, full survival), as indicated by the color bar.

Evaluated on the NCI-ALMANAC dataset^35^ comprising 1,807,365 dose-dependent viability measurements across 41 cell lines and 105 compounds, XACT-Dual maintained high predictive fidelity across the entire dynamic range of combinatorial cell viability responses, from near-complete cell death to full survival (**Fig. 3B**). Even in the rigorous leave-cells-out scenario, the model effectively suppressed prediction error, demonstrating robustness to cellular heterogeneity and generalization to unseen biological contexts (**Fig. 3C**). Benchmarking against ComboLTR^36^ and state-of-the-art DD-PRiSM^37^ confirmed that XACT-Dual outperformed all competitors across every evaluated metric under both random split cross-validation (**Fig. 3D, Supplementary Table 4**). To further evaluate generalizability under the more stringent leave-cells-out condition, we compared XACT-Dual with the best-performing competitor, DD-PRiSM. XACT-Dual achieved statistically significant improvements across all evaluated metrics with relative improvements of up to 20% (\*\**p* < 0.01; **Fig. 3E**, **Supplementary Table 5**), confirming that XACT-Dual captures generalizable combinatorial pharmacodynamic rules rather than overfitting to the statistical regularities of the training distribution.

A defining capability of XACT-Dual is its reconstruction of the full two-dimensional dose-response manifold for any drug pair and cell line combination, rather than predicting a single synergy scalar. This was confirmed by three-dimensional surface analysis of representative combinations, in which XACT-Dual faithfully reproduced the experimentally observed dose-response surfaces including the characteristic synergy profiles of oxaliplatin plus fluorouracil in HCT-116 colorectal cancer cells^35^ (**Fig. 3F**), as well as additional drug pairs spanning distinct synergy patterns across multiple cancer cells (**Supplementary Fig. 3**).

### XACT-Clinical resistance scores independently stratify overall survival in the TCGA pan-cancer cohort

Having established the predictive accuracy of XACT-Dual in preclinical benchmarks, we next sought to evaluate whether the pharmacological representations learned from cell line data retain sufficient biological signal to stratify clinical treatment outcomes in real patients. To adapt the XACT framework to this task, the final regression layer of XACT-Dual was replaced with a binary classification head and optimized on The Cancer Genome Atlas (TCGA)^38^ treatment outcome data, yielding XACT-Clinical. Clinical responses were defined according to RECIST criteria^39^, with Complete Response (CR) or Partial Response (PR) designated as responders and those exhibiting Stable Disease (SD) or Progressive Disease (PD) designated as non-responders. The model was adapted under a strict ten-fold patient-level cross-validation scheme. The resulting output, designated as the XACT resistance score, represents the predicted probability of non-response for a given patient transcriptome, drug pair, and maximum plasma concentration (*C*_max_) combination (**Fig. 4A**). All XACT resistance scores used for clinical outcome and survival analyses were out-of-fold predictions generated when a patient was assigned to the test fold, thereby completely preventing data leakage arising from within-patient correlation.

**Fig. 4.**
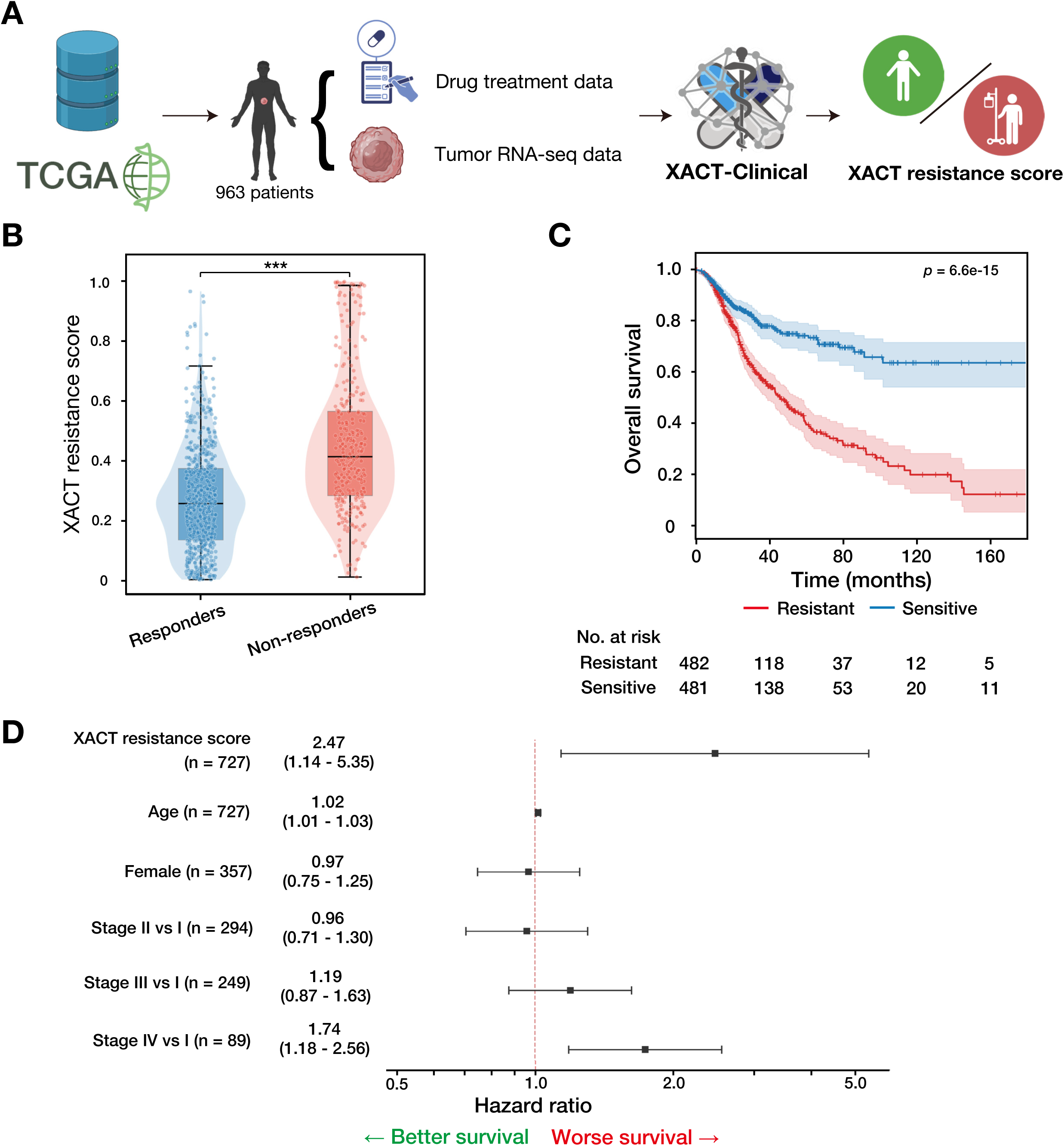
Prospective *in silico* clinical trial validation and independent prognostic utility of XACT-Clinical in the TCGA pan-cancer cohort. **(A)** *In silico* clinical trial protocol. Tumor RNA-seq data and combination chemotherapy records from 963 patients in the TCGA pan-cancer cohort were used to evaluate the clinical translational utility of XACT-Clinical. The final regression layer of XACT-Dual was replaced with a binary classification head and optimized on TCGA treatment outcome data to yield XACT-Clinical. Clinical response labels were defined according to RECIST criteria, designating patients achieving Complete Response (CR) or Partial Response (PR) as responders and those exhibiting Stable Disease (SD) or Progressive Disease (PD) as non-responders. The resulting output, designated as the XACT resistance score, represents the predicted probability of non-response for a given patient transcriptome, drug pair, and the corresponding maximum plasma concentration values. **(B)** Discrimination of clinical responders and non-responders by XACT resistance score. Violin plots displaying the distribution of XACT resistance scores stratified by RECIST-defined clinical outcome. Responders received significantly lower XACT resistance scores than non-responders (Mann-Whitney U test, \*\*\**p* < 0.0001). **(C)** Prognostic stratification via Kaplan-Meier survival analysis. Patients were stratified into predicted-resistant (red) and predicted-sensitive (blue) groups at the median XACT resistance score, and overall survival probability was estimated using the Kaplan-Meier method. Shaded regions indicate 95% confidence intervals. Numbers at risk are shown below the survival curves. **(D)** Independence of the XACT resistance score as a prognostic factor. Forest plot from multivariate Cox proportional hazards regression incorporating the XACT resistance score alongside established clinical covariates including tumor stage, patient age, gender, and cancer type. Hazard ratios (HR) and 95% confidence intervals are shown for each covariate. After full adjustment for all clinical covariates, the XACT resistance score remained the strongest independent prognostic factor (HR = 2.47, 95% CI: 1.14–5.35), with a point estimate exceeding that of advanced tumor stage (Stage IV vs. Stage I: HR = 1.74, 95% CI: 1.18–2.56), demonstrating that XACT-Clinical captures intrinsic molecular determinants of therapeutic vulnerability that are orthogonal to standard pathological staging.

Patients who achieved an objective clinical response to their prescribed regimen received significantly lower XACT resistance scores than non-responders (Mann-Whitney U test, *p* < 0.001, **Fig. 4B**), demonstrating that XACT resistance scores are significantly associated with objective clinical treatment outcomes across diverse cancer types and combination regimens. We further evaluated whether XACT resistance scores stratify long-term survival by dichotomizing patients at the median resistance score into predicted-sensitive and predicted-resistant groups and estimating overall survival by the Kaplan-Meier method. The resulting survival curves revealed a striking and durable separation between groups throughout the follow-up period (log-rank *p* = 6.6 × 10^-15^; **Fig. 4C**), with patients in the predicted-sensitive group exhibiting markedly prolonged overall survival relative to those in the predicted-resistant group.

To verify that this survival advantage reflected intrinsic pharmacogenomic vulnerability rather than confounding clinical characteristics, we performed multivariate Cox proportional hazards regression^40^ incorporating tumor stage, patient age, gender, and cancer type as covariates (**Fig. 4D**, **Supplementary Table 6**). After full adjustment for all clinical covariates, the XACT resistance score emerged as the strongest independent prognostic factor with a Hazard Ratio (HR) of 2.47 (95% CI: 1.14–5.35, *p* = 0.022), with a point estimate exceeding that of advanced tumor stage (Stage IV vs. Stage I: HR = 1.74, 95% CI: 1.18–2.56). These results establish that XACT-Clinical captures intrinsic molecular determinants of therapeutic vulnerability that are orthogonal to and not explained by standard pathological staging, providing compelling evidence for its utility as a clinically translatable independent prognostic tool.

### XACT-Clinical reveals cancer-type-specific pharmacogenomic vulnerabilities and nominates alternative therapeutic regimens

Sarcoma and pancreatic adenocarcinoma represent two of the most therapeutically refractory solid malignancies, with limited second-line options and poor outcomes for patients who fail standard-of-care regimens. We applied XACT-Clinical to conduct systematic virtual screening analyses in both cancer types, seeking to identify computationally superior alternative regimens for patients predicted to be non-responsive to their prescribed treatment.

XACT resistance scores successfully stratified patients by clinical outcome, with non-responders exhibiting significantly higher predicted resistance than responders to their prescribed regimen in the TCGA sarcoma (TCGA-SARC) cohort (**Fig. 5A**) and a consistent trend observed in the pancreatic adenocarcinoma (TCGA-PAAD) cohort (**Supplementary Fig. 4A**), confirming the capacity of XACT-Clinical to reflect differential drug sensitivity across patient subgroups in histologically distinct malignancies.

**Fig. 5.**
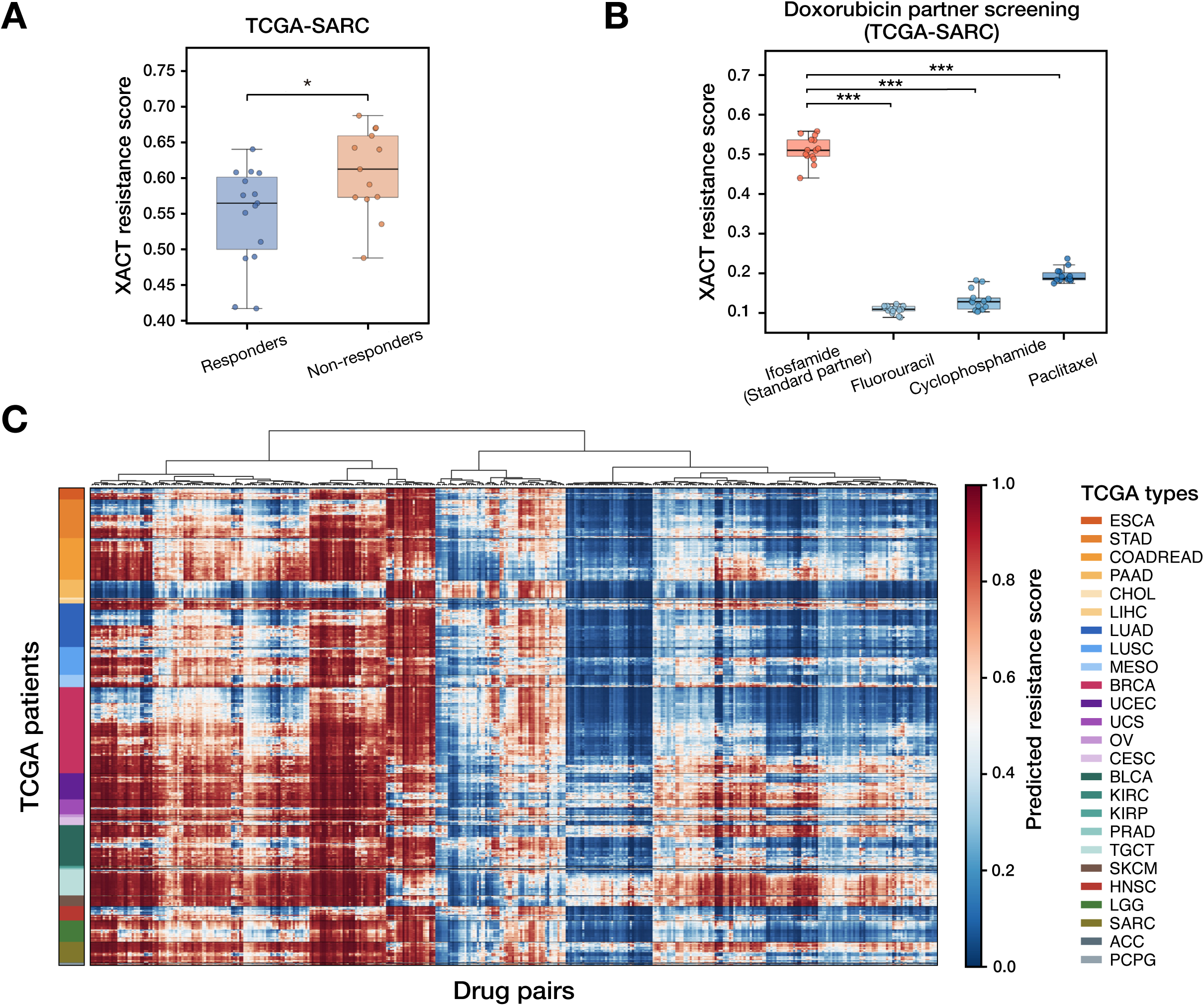
XACT-Clinical reveals cancer-type-specific pharmacogenomic vulnerabilities and nominates alternative therapeutic regimens. **(A)** Recapitulation of clinical outcomes in sarcoma. Box plots displaying XACT resistance scores for TCGA sarcoma (TCGA-SARC) patients stratified by RECIST-defined clinical outcome. Non-responders (SD/PD) exhibited significantly higher resistance scores than responders (CR/PR) (\**p* < 0.05), demonstrating that XACT resistance scores reflect the differential drug sensitivity underlying observed clinical outcomes. **(B)** *In silico* doxorubicin anchor-partner screening for sarcoma. Box plots displaying XACT resistance scores for the three highest-ranked alternative combination partners for doxorubicin across non-responding SARC patients, with the standard-of-care partner ifosfamide shown for reference. Statistical significance of pairwise comparisons against ifosfamide was assessed by two-sided *t*-test (\*\*\**p* < 0.001). **(C)** Pan-cancer pharmacogenomic resistance landscape. Heatmap displaying XACT resistance scores for all 963 TCGA patients across 410 drug pairs which exhibit high XACT resistance score variance between cancer types. Patients are arranged along the vertical axis sorted by cancer type as indicated by the color-coded legend, and drug pairs are arranged along the horizontal axis. The structured heterogeneity of the resistance landscape across cancer types and drug pairs reveals substantial inter-tumoral variability in predicted sensitivity, providing a comprehensive atlas of combinatorial pharmacogenomic vulnerability across the TCGA pan-cancer cohort.

*In silico* anchor-partner screening identified superior combination partners within established cytotoxic backbone therapies in both cancer types. With doxorubicin fixed as the cytotoxic anchor in SARC, fluorouracil and cyclophosphamide were identified as superior alternatives to the standard-of-care partner ifosfamide^41,42^, yielding nearly five-fold reductions in predicted resistance with markedly tighter variance across the non-responding cohort (*p* < 0.001; **Fig. 5B**), suggesting that substitution with agents inducing orthogonal mechanisms of DNA damage may overcome the resistance barriers associated with ifosfamide-based therapy. With gemcitabine fixed as the anchor in PAAD, capecitabine, cisplatin, and sorafenib were identified as the highest-ranked combination partners, all demonstrating significantly lower predicted resistance than the standard-of-care partner paclitaxel^43^ (**Supplementary Fig. 4B**). The prioritization of sorafenib is of particular biological interest, as prior clinical trials of this multi-kinase inhibitor in combination with gemcitabine have not demonstrated improved overall survival in unselected PAAD patients^44^, suggesting that transcriptomic stratification by XACT-Clinical may identify the subpopulation most likely to benefit from this combination. Furthermore, virtual first-line screening nominated alternative regimens exhibiting substantially lower predicted resistance than the actual prescribed treatments across the non-responding subgroup in both cancer types. In SARC and PAAD, three alternative regimens were computationally prioritized (**Supplementary Figs. 4C and 4D**).

To provide a comprehensive pan-cancer view of combinatorial pharmacogenomic vulnerability, XACT resistance scores were computed across all 963 TCGA patients and 410 drug pairs and assembled into a pan-cancer resistance landscape (**Fig. 5C**). While predicted resistance profiles differed broadly across cancer types, with mesothelioma and sarcoma exhibiting broadly elevated predicted resistance and testicular germ cell tumors and breast cancer displaying extensive regions of predicted sensitivity, hierarchical clustering within each cancer type revealed substantial inter-patient heterogeneity in pharmacogenomic sensitivity, demonstrating that patients sharing the same diagnosis can exhibit markedly distinct vulnerability profiles to the same combination regimen. This intra-cancer-type heterogeneity, driven by differences in underlying tumor transcriptomes rather than histological classification alone, underscores the necessity of patient-level pharmacogenomic profiling and provides the rational foundation for individualized therapeutic selection in precision oncology. Concurrent clustering of drug pairs identified structured groupings of combinations with shared activity spectra, suggesting that therapeutic vulnerability is organized according to convergent mechanisms of action rather than individual drug identity. This atlas provides a systematic, data-driven framework for identifying cancer types and drug pair contexts in which the greatest potential for therapeutic improvement through computational repurposing exists, and establishes a prioritized catalogue of combination strategies warranting prospective experimental validation.

### MIL-augmented XACT-Clinical extends prognostic stratification and therapeutic nomination to complex multi-drug regimens

Although the majority of standard oncological regimens involve three or more agents, existing pairwise prediction frameworks^15,16,36,37^ are inherently unable to capture the emergent pharmacological effects that arise from higher-order drug interactions, and no computational model has yet been developed to address this gap. To address this fundamental limitation, XACT-Clinical was extended by incorporating a multi-instance learning (MIL) framework^45^ that enables holistic evaluation of multi-drug combinations without decomposing complex regimens into independent pairwise interactions whose sum fails to represent the true combinatorial effect (**Fig. 6A**).

**Fig. 6.**
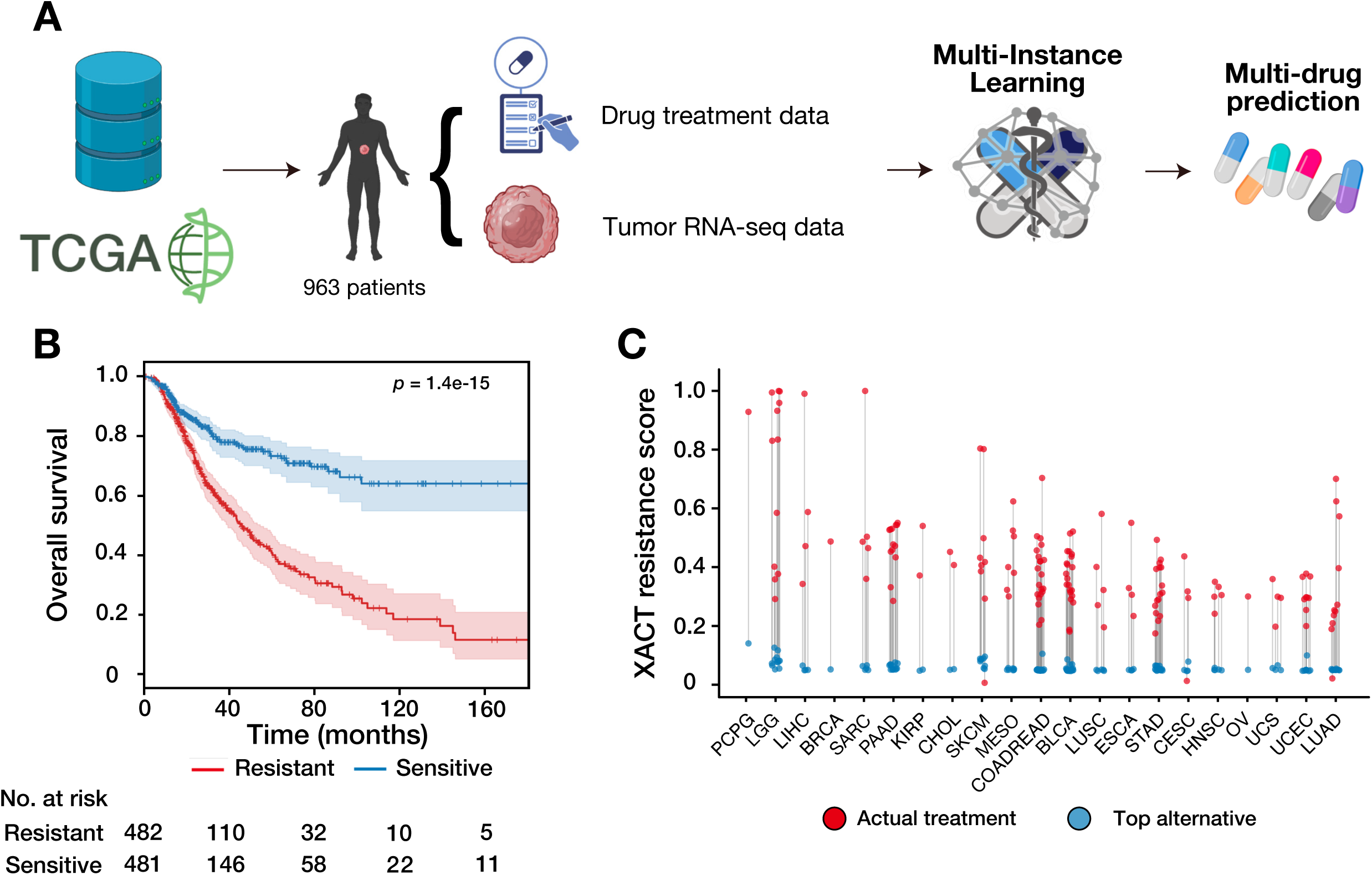
Extension to multi-drug prediction via multi-instance learning and pan-cancer therapeutic alternative nomination. **(A)** Multi-instance learning framework for multi-drug prediction. Schematic illustrating the extension of XACT-Clinical beyond pairwise combinations to regimens comprising three or more agents using a multi-instance learning (MIL) architecture. Rather than decomposing multi-drug regimens into independent pairwise interactions, the MIL framework evaluates the complete drug combination holistically by treating each patient as a bag of drug pair instances and aggregating instance-level resistance scores through min-pooling, enabling the model to capture emergent pharmacological effects not reducible to the sum of pairwise interactions. **(B)** Prognostic stratification by MIL-augmented XACT-Clinical resistance scores. Kaplan-Meier survival curve for the TCGA pan-cancer cohort stratified into predicted-resistant and predicted-sensitive groups at the median MIL-derived resistance score. Shaded areas represent 95% confidence intervals. Patients in the predicted-sensitive group exhibited markedly prolonged overall survival relative to those in the predicted-resistant group (log-rank *p* = 1.4 × 10^-15^), demonstrating that the MIL framework preserves and extends the prognostic stratification capability established for pairwise predictions. Numbers at risk are shown below the survival curves. **(C)** Pan-cancer landscape of predicted therapeutic vulnerability and computationally nominated three-drug alternatives. Lollipop plot displaying the XACT resistance score for each patient’s actual prescribed treatment regimen (red) and the top-ranked alternative drug triplet identified by virtual screening (blue) for each TCGA cancer type. Each red point represents the predicted resistance of the standard-of-care combination for an individual patient, and each blue point represents the lowest predicted resistance achievable by substituting one agent with a computationally nominated alternative. The systematic displacement of blue points toward lower predicted resistance values relative to red points across the majority of cancer types indicates that computationally nominated triplet regimens consistently exhibit lower predicted resistance than the standard-of-care combination, identifying cancer types with the greatest potential for therapeutic improvement through computational drug repurposing.

Applied to the full TCGA cohort of 963 patients, the MIL-augmented XACT-Clinical generated predicted resistance scores that successfully stratified patient survival with a magnitude comparable to that observed for pairwise predictions (log-rank *p* = 1.4 × 10^-15^; **Fig. 6B**), demonstrating that holistic modeling of multi-drug regimens preserves and extends the prognostic signal captured by the pairwise framework. When applied to screen computationally nominated three-drug combinations across each cancer type in the TCGA cohort, MIL-augmented XACT-Clinical identified triplet regimens exhibiting substantially lower predicted resistance than the corresponding standard-of-care drug pairs in the majority of cancer types examined (**Fig. 6C**). The systematic displacement of predicted resistance scores toward lower values for computationally nominated triplets relative to prescribed treatments identifies cancer types in which the incremental addition of a rationally selected third agent may overcome residual resistance barriers that are insurmountable with dual-drug therapy alone, positioning MIL-augmented XACT-Clinical as a scalable computational strategy for the rational design of higher-order therapeutic regimens across the full spectrum of oncological practice.

## Discussion

This study establishes XACT as a multi-stage deep learning framework capable of predicting dose-dependent combinatorial drug responses using transcriptomic profiles and chemical structure alone, and demonstrates that the pharmacological rules encoded from preclinical data are sufficiently generalizable to stratify clinical treatment outcomes in real patients. The persistent translational gap between pharmacogenomics and clinical practice has historically arisen from three compounding failures. Most existing models reduce drug sensitivity to a single scalar metric, sacrificing the dose-dependent resolution necessary for rational therapeutic optimization. Those that achieve higher predictive resolution typically rely on multi-omics data modalities unavailable in routine clinical settings, fundamentally restricting real-world applicability. Finally, even the most accurate preclinical models have rarely been validated against clinical outcomes, leaving their prognostic utility undemonstrated. XACT addresses all three failures within a unified framework, predicting full dose-dependent combinatorial drug responses from clinically accessible transcriptomic profiles alone and demonstrating that these predictions stratify patient survival as an independent prognostic factor superior to standard clinical staging.

The superior generalizability of XACT-Single and XACT-Dual stems from architectural inductive biases that explicitly encode pharmacological reality rather than relying on implicit feature memorization. Conventional drug response models treat drug-cell interactions as static feature pairs, forcing the network to memorize context-specific associations that do not transfer across biological systems. The X-Linear Attention mechanism departs from this paradigm by computing second-order bilinear interactions between molecular drug substructures and intracellular pathway activities, enabling the model to learn the generalized rules of pharmacological context-dependency rather than specific drug-cell combinations. The asymmetric cross-branch attention of XACT-Dual extends this principle to the combinatorial domain by explicitly encoding the directional modulation of one drug’s mechanism of action by its combination partner, mathematically formalizing the biological scenario in which one agent remodels the cellular signaling landscape and thereby conditions the environment in which its partner operates. This architectural design is directly responsible for the capacity to generalize to unseen transcriptomic landscapes, as demonstrated by the less than 3% degradation in predictive accuracy between random split and leave-cells-out evaluations.

A distinctive feature of XACT that separates it from virtually all existing drug response prediction frameworks is its capacity to reconstruct full dose-dependent cell viability curves rather than reducing therapeutic efficacy to a scalar IC₅₀ or synergy score. This capability is not merely a technical refinement but a clinically essential advance: rational dose optimization, the identification of therapeutic windows, and the prediction of concentration-dependent synergistic effects all require knowledge of the complete pharmacodynamic trajectory. The prospective experimental validation in SaOS-2 osteosarcoma cells, a cancer type entirely absent from the training data, further demonstrates that XACT encodes transferable pharmacological rules rather than overfitting to the statistical patterns of the training distribution. The mechanistic interpretability analyses provide an additional and independent line of evidence that the model has internalized biologically valid pharmacological relationships. The autonomous identification of KRAS and RAF signaling as dominant transcriptomic determinants of trametinib sensitivity, and the prioritization of the allosteric pharmacophore as the structural driver of predicted potency, recapitulate canonical MEK inhibitor pharmacology without explicit biological supervision (Supplementary Fig. 1), confirming that XACT predictions are grounded in causal biological mechanisms rather than dataset-specific statistical artifacts.

The most transformative contribution of this work lies in the demonstrated clinical utility of XACT-Clinical. The XACT resistance score emerged as the strongest independent prognostic factor in multivariate Cox regression adjusted for tumor stage, patient age, sex, and cancer type (HR = 2.47, 95% CI: 1.14–5.35, *p* = 0.022), with a point estimate exceeding that of Stage IV metastasis. This suggests that XACT captures a dimension of intrinsic molecular drug responsiveness that is orthogonal to and not encoded by standard pathological variables, offering a fundamentally new axis of prognostic information for precision oncology. The virtual screening analyses in sarcoma and pancreatic adenocarcinoma further illustrate the practical utility of this framework, nominating biologically plausible alternative regimens supported by established clinical and mechanistic evidence and providing a prioritized catalogue of combination strategies warranting prospective experimental evaluation. The extension to higher-order regimens via MIL-augmented XACT-Clinical demonstrates that the prognostic stratification capability established for pairwise predictions is preserved and extended to multi-drug combinations, broadening the clinical applicability of the framework to the complex polypharmacy regimens that characterize modern oncological practice.

Despite these advances, several limitations warrant consideration. The clinical validation relied on retrospective bulk RNA-seq data and binary RECIST classifications, which may not fully capture the continuous dynamics of tumor response or the spatial and cellular heterogeneity of human tumors. Single-cell and spatial transcriptomic data, as they become more broadly available in clinical settings, could substantially enhance the resolution of cellular state encoding in future iterations of the framework. Furthermore, XACT-Clinical does not currently account for patient-specific pharmacokinetics or intratumoral drug distribution, both of which influence the effective concentration experienced by tumor cells *in vivo*. Integrating pharmacokinetic-pharmacodynamic modeling into the XACT framework represents a necessary next step toward refining dose optimization for individual patients. Finally, although the MIL framework extends combinatorial prediction to regimens of three or more agents, the explicit modeling of higher-order drug interactions through generalized multi-drug attention mechanisms remains an important direction for future architectural development, particularly for the rational design of complex regimens such as FOLFOX that underpin standard oncological care.

In conclusion, XACT demonstrates that biologically grounded deep learning can successfully traverse the translational divide by decoding the fundamental rules of drug sensitivity from transcriptomic and structural data. The hierarchical progression from monotherapy response prediction through combinatorial modeling to clinical prognostication, and ultimately to prospective experimental validation and pan-cancer therapeutic repurposing, establishes XACT as a scalable, interpretable, and translation-ready framework that advances precision oncology from computational prediction toward data-driven therapeutic prescription.

## Materials and Methods

### Data acquisition and preprocessing

#### Cancer cell line drug response datasets

Large-scale drug sensitivity data were sourced from two primary repositories. IC_50_ values and dose-dependent cell viability measurements for monotherapy response prediction were obtained from the Genomics of Drug Sensitivity in Cancer (GDSC) database^8^, comprising 118,896 IC₅₀ measurements across 959 cancer cell lines and 162 compounds as well as 1,509,214 dose-dependent cell viability measurements across 941 cancer cell lines and 228 compounds. Combinatorial drug response data were obtained from the NCI-ALMANAC dataset^35^, which provides dose-dependent cell viability measurements for approximately 5,000 drug pairs across 60 human tumor cell lines.

#### Transcriptomic feature engineering

Raw RNA-seq data in FPKM units for all cell lines were retrieved from the GDSC database. To ensure robust, biologically interpretable input features, gene-level expression values were log-transformed according to x′=log_2_(x+1) and subsequently transformed into pathway-level activity scores through Single Sample Gene Set Enrichment Analysis (ssGSEA)^22,46^, computed using the GSEApy library^47^. Gene sets were drawn from two complementary collections catalogued in the Molecular Signatures Database (MSigDB)^48^: the 50 Hallmark gene sets^49^, which capture fundamental and well-defined biological processes, and 189 C6 oncogenic signature gene sets, which capture cancer-specific transcriptional dysregulation. This pipeline yielded a fixed-length feature vector of 239 dimensions for each cancer cell line.

#### Multi-modal chemical representation

To capture both the explicit topological structure and the latent bioactivity profiles of small molecules, a dual-modality embedding strategy was employed that integrates graph-based representations encoding local atomic interactions with pre-computed bioactivity signatures encoding global physicochemical and biological contexts. Canonical SMILES strings for all compounds in the drug response datasets were obtained through PubChem^50^, using compound names as search queries.

To encode the broader biological and physicochemical context of each drug, the Chemical Checker (CC)^21^ was employed. The CC framework harmonizes heterogeneous bioactivity data into compact 128-dimensional vector signatures, providing a unified representational space for small molecules. The five Level A signatures (A1–A5) were incorporated, which collectively encode complementary aspects of molecular identity: A1 encodes standard two-dimensional topological fingerprints, A2 captures three-dimensional structural descriptors, A3 encodes Murcko scaffold representations^51^, A4 represents predefined structural substructure keys, and A5 encodes global physicochemical properties including molecular weight and logP. These signatures were selected for their capacity to facilitate the inference of missing bioactivity data and their seamless compatibility with deep learning architectures.

To explicitly model molecular topology and local atomic neighborhoods, molecular graphs *G* = (*V*, *E*) were constructed where *V* represents the set of atoms and *E* denotes the set of chemical bonds. Graph construction and feature extraction were implemented using the RDKit cheminformatics library (version 2024.03.6) and NetworkX (version 3.4.2). Following existing work^12^, each atom node *V* was initialized with a 78-dimensional feature vector constructed via one-hot encoding of five atomic properties. The five atomic property-derived encoded node features comprised the atom symbol encoded across 44 allowable atomic types encompassing all atomic species present in the evaluated drug libraries, the atom degree representing the number of covalent bonds ranging from 0 to 10 to accommodate the full valency range observed in the evaluated drug libraries, the total number of hydrogens encompassing both implicit and explicit species over the same range, the implicit valence over the same range, and a binary aromaticity flag indicating membership in an aromatic ring system. The concatenated feature vector for each atom was normalized by the sum of its elements to ensure numerical stability during training. Molecular edges were encoded as a directed graph in Coordinate Format (COO) to ensure compatibility with the PyTorch Geometric^52^ framework.

### XACT-Single architecture

The XACT-Single monotherapy response prediction module was designed to predict the sensitivity of individual drugs against specific cancer cells, yielding both IC_50_ estimates and continuous dose-dependent cell viability profiles. Conceptually, the model frames the interaction between a drug and a cell line as a query-key attention problem of the transformer architecture^53^, in which the cellular state serves as a dynamic query that selectively retrieves the most task-relevant features from the drug’s multi-modal profile.

#### Chemical feature encoding

To construct a richly informative drug representation, the XACT-Single encoder maintains a structured multi-view representation that preserves the distinct information content of each modality. Drug features were processed through two parallel branches designed to capture complementary aspects of molecular identity.

The structural branch processed each molecular graph through a 3-layer Graph Isomorphism Network (GIN)^20^, in which each layer consists of a Multi-Layer Perceptron (MLP) followed by Batch Normalization and ReLU^54^ activation. Global add pooling was applied to aggregate node-level features into a graph-level representation of 256 dimensions, as formulated below:

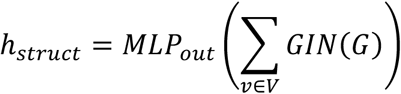

The hidden dimension was set to 128 with a dropout rate of 0.2.

The bioactivity branch processed the five Chemical Checker signatures (A1–A5) through five independent parallel feed-forward networks (FFNs), each projecting its respective signature into a shared latent space of 256 dimensions. For each view *i*, the two-layer FFN is defined as:

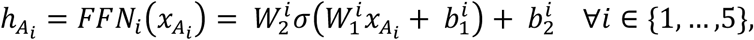

where 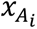 is the input feature vector for the *i*-th Chemical Checker signature, 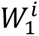 and 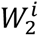 are learnable weight matrices, 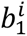 and 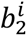 are biases, and σ represents the ReLU activation function. Rather than concatenating the resulting feature vectors, the six view embeddings — comprising the structural descriptor and the five bioactivity signatures —were stacked along a dedicated view axis to form a structured sequence:

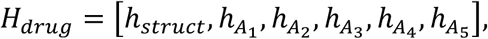

yielding a tensor of shape (*B*, *N*_*views*_, *d*_*model*_), where *N*_*views*_ = 6. This representation preserves the semantic boundaries between modalities and enables the subsequent attention mechanism to dynamically weight individual drug views conditioned on the cellular transcriptomic context.

#### Cell line feature encoding

Pathway-level activity scores derived from ssGSEA^22^ were used as input features. These 239-dimensional normalized enrichment score (nes) vectors were projected into the shared latent space of 256 dimensions through a 2-layer MLP, ensuring representational compatibility with the multi-view drug embeddings:

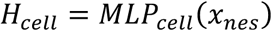

The resulting cellular embedding *H*_*cell*_ serves as the query signal in the subsequent attention mechanism, encoding which aspects of the drug’s multi-modal profile are most pharmacologically relevant given the transcriptomic state of the target cell.

#### X-Linear Attention block

The X-Linear Attention mechanism^23^ was adapted to model the complex, non-linear interactions between the cellular state and the multi-modal drug representation. Standard attention mechanisms, whether additive or dot-product^53^, fuse the query and key through linear operations to model exclusively first-order interactions. X-Linear Attention overcomes this limitation by employing bilinear pooling, which explicitly captures second-order feature interactions by computing the outer product between projected query and key representations, enabling the model to detect synergistic co-activation patterns between molecular substructures and intracellular signaling pathways that would remain invisible to linear fusion strategies.

Within this framework, the cell line embedding *H*_*cell*_ serves as the query G, encoding the cellular context that determines which drug features are pharmacologically salient, while the multi-view drug embeddings *H*_*dru*g_ serve as both the Key *K* and Value *V*. This asymmetric query-key assignment reflects the biological premise that the transcriptomic state of the cell defines the functional landscape against which drug features are evaluated.

#### Low-rank bilinear pooling

Although full bilinear pooling provides an expressive mechanism for capturing second-order feature interactions^55^, its direct computation yields a feature space whose dimensionality grows as the product of the input dimensions, rendering it computationally prohibitive at pharmacogenomic scales. The X-Linear Attention (X-LAN) module therefore employs low-rank bilinear pooling^56^, which approximates the outer product through element-wise multiplication in a projected latent space. For each drug view, the joint bilinear representation *B*^*k*^ between the cellular query G and the

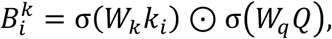

where *σ* denotes the ReLU activation, ⊙ represents element-wise multiplication, and *W*_*k*_ and *W*_*q*_ are learnable projection matrices that independently compress the key and query representations into a shared low-dimensional interaction space.

#### Dual attention distribution

The X-Linear Attention mechanism operates simultaneously across two complementary axes of selectivity through spatial attention over drug views and channel-wise attention over feature dimensions. Spatial attention derives a scalar relevance score for each drug view through a learned projection:

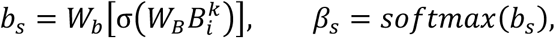

where *β*_*s*_ encodes a probability distribution over the *N* drug views reflecting the relative pharmacological salience of each modality. Channel-wise attention is achieved through a Squeeze-and-Excitation operation^57^, in which bilinear representations are compressed into a global channel descriptor *B̅* via view-wise average pooling and then passed through a sigmoid-gated excitation layer:

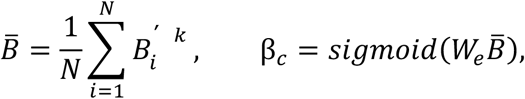

where *β*_*c*_ acts as a soft feature gate, rescaling each latent dimension according to its estimated global relevance. The final attended representation is obtained by aggregating the bilinear value representations weighted by spatial attention and modulated by the channel-wise gate:

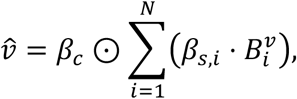

where 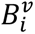 denotes the bilinear interaction between the cellular query and the *i*-th drug value embedding, computed through the same low-rank factorization. The attended representation *v̂* constitutes a compact, second-order encoding of the drug-cell interaction that jointly reflects the molecular substructure most pertinent to the cellular signaling context.

#### Regression module

A three-layer non-linear regression module projects *v̂* onto the continuous log-IC_50_ response scale through an expand-compress architecture that first lifts the representation into a higher-dimensional space to disentangle the latent factors encoded by the attention mechanism, then compresses it into a compact bottleneck representation. ReLU activation and dropout regularization (rate = 0.3) were applied after each of the first two transformations:

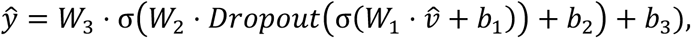

where *ŷ* represents the predicted log-normalized IC_50_ value and *W*_*l*_, *b*_*l*_ denote the weight matrix and bias vector of the *l*-th layer, respectively.

#### Cell viability prediction

Whereas the IC_50_ module outputs a single sensitivity scalar per drug-cell pair, predicting dose-dependent cell viability requires modeling a continuous survival fraction across the full concentration ranges. The feature extraction pipeline remains architecturally identical to the IC_50_ model, but the regression module was redesigned to condition the survival prediction on drug concentration through a late-fusion strategy. The global interaction vector *v̂* is first transformed by a dense layer into a high-level biological feature representation ℎ_*bio*_ that encodes the intrinsic sensitivity of the cell to the drug independently of concentration. The log-transformed drug concentration *c* is subsequently concatenated with ℎ_*bio*_, ensuring that dosage modulates the prediction only after the drug-cell biological interaction has been fully encoded. The concatenated representation is then passed through a final MLP to predict the fractional cell viability:

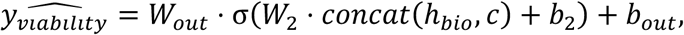

where *c* denotes the log-transformed drug concentration and all other notation is consistent with the regression module above.

#### Hyperparameter scaling

Modeling the full dose-response curve demands substantially greater representational capacity than predicting a single IC_50_ scalar. To reflect this increased complexity, the dimension of the joint bilinear representation was quadrupled from 256 to 1024, and the intermediate hidden dimensions of both drug and cell embedding networks were correspondingly increased from 128 to 512.

#### Training strategy and hyperparameters

All XACT-Single models were optimized by minimizing the Mean Squared Error (MSE) between predicted and observed responses using the AdamW optimizer^58^ with a weight decay of 0.001. The learning rate was initialized at 1×10^-4^ and reduced by a factor of 0.8 every three epochs through a step-decay schedule. Training was conducted for a maximum of 100 epochs with a mini-batch size of 128, with the checkpoint attaining the lowest MSE on the held-out validation retained as the final model. The framework was implemented in Python using PyTorch^59^ and PyTorch Geometric, with all experiments performed on a single NVIDIA H100 GPU on the TSUBAME supercomputer.

#### Dose-response curve prediction and fitting

To evaluate dose-dependent pharmacodynamic predictions in biologically interpretable terms, models trained under 10-fold leave-cells-out cross-validation were applied to held-out cell lines to generate viability predictions across the tested concentration range. For each curated drug-cell line pair, both experimentally observed and the model-predicted viability values were plotted against drug concentrations on a logarithmic scale and fitted to a four-parameter Hill equation:

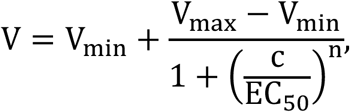

where *V* represents fractional cell viability, *c* is the drug concentration, *V*_*mbx*_and *V*_*min*_are the upper and lower asymptotic viability values, *EC*_50_ is the half-maximal effective concentration, and *n* is the Hill coefficient governing the steepness of the sigmoidal curve. The curve fitting was performed using non-linear least squares optimization, with *V*_*mbx*_and *V*_*min*_ constrained between 0 and 2.0 to ensure biologically plausible parameter estimates.

### XACT-Dual architecture

To predict the dose-dependent viability of cells exposed to drug pairs, the XACT-Single architecture was extended into a weight-shared Siamese network framework augmented with asymmetric cross-branch attention. The monotherapy encoders were held fixed as stable feature extractors, ensuring that the fundamental pharmacological representations of each drug are preserved and preventing overfitting to the comparatively sparse combinatorial training data. Each branch independently processes the features of one drug alongside the shared cell line representation through the full XACT-Single pipeline to yield a global interaction vector ℎ_*k*_ and a multi-view attention map *α*_*k*_. Formally, for branch *k*:

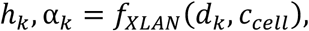

where *f*_*XLAN*_ represents the frozen XACT-Single encoder, *d*_*k*_ the molecular representation of drug *k*, and *c*_*cell*_ the shared transcriptomic representation of the target cell.

#### Cross-branch attention fusion

Two trainable X-Linear Attention blocks were introduced to perform cross-branch reasoning across the parallel drug encoders. These fusion blocks perform inter-drug attention in which the global representation of one drug serves as the query that interrogates the multi-view attention map of its combination partner, explicitly modeling the biological scenario in which one agent remodels the intracellular signaling landscape and thereby conditions the environment in which its partner operates. Specifically, the first fusion block uses the global interaction vector of drug A (ℎ_*A*_) as the query to attend over the multi-view attention map of drug B (*α*_*B*_):

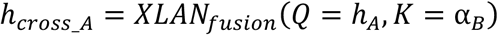

The second fusion block applies the symmetric operation using ℎ_*B*_ as the query to attend over *α*_*A*_, yielding ℎ_*cross*_*B*_.

#### Combination Regressor

The cross-attended representations ℎ_*cross*_*A*_ and ℎ_*cross*_*B*_ were concatenated to form a unified combination embedding. The log-transformed concentrations of both drugs were appended to this embedding through the same late-fusion strategy employed in the monotherapy viability model, and the concatenated representation was passed through a three-layer MLP that predicts the two-drug cell viability:

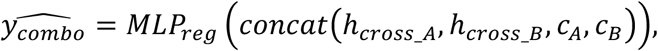

where *c*_*A*_ and *c*_*B*_ denote the log-transformed concentrations of drugs A and B, respectively.

#### Transfer learning and training strategy

The weights of the drug embedding module, cell embedding module, and primary X-Linear Attention blocks were initialized from the XACT-Single viability model and frozen throughout combinatorial training, confining the parameter search to the cross-branch fusion blocks and the combination regressor where combinatorial training signal is most informative. The model was trained on the NCI-ALMANAC dataset using the AdamW optimizer with a learning rate of 1×10^-4^ and a weight decay of 2× 10^-4^, minimizing the Mean Squared Error (MSE) between the predicted and observed two-drug cell viability.

### Model benchmarking

#### Implementation and benchmarking of existing monotherapy IC_50_ prediction models

To rigorously evaluate the predictive performance of XACT-Single, benchmarking was performed against three established pathway-guided deep learning models designed for monotherapy IC₅₀ prediction: PathDNN^11^, ConsDeepSignaling (CDS)^10^, and HiDRA^13^, together with a standard multi-layer perceptron (MLP) baseline. All baseline models were executed using standardized re-implementations provided in a dedicated benchmarking repository^60^ under a controlled computational environment comprising Python (version 3.9.7), PyTorch (version 1.11.0), pandas (version 1.3.4), numpy (version 1.20.3), scipy (version 1.7.1), and scikit-learn (version 0.24.2). Hyperparameter tuning was strictly governed by the optimized configurations established in the repository for each validation strategy. Model evaluation was conducted under 5-fold cross-validation applied to both random split and leave-cells-out conditions.

#### Dose-response surface reconstruction

To evaluate the fidelity of combinatorial viability predictions against experimentally observed pharmacological landscapes, dose-response surfaces were reconstructed for clinically established first-line combination therapies. For each evaluated cell line, predictions were generated using the XACT-Dual model from the corresponding leave-cells-out cross-validation fold, ensuring all predictions were derived from strictly out-of-sample data. A dense 25×25 concentration spanning the minimum and maximum experimental concentrations of both drugs was defined, and model inference was performed across this grid to yield a continuous high-density predicted viability surface. Experimental viability matrices were reconstructed into corresponding three-dimensional surfaces by smoothing and mapping sparse empirical measurements to the 25×25 computational grid utilizing linear regular grid interpolation. Predicted and experimental surfaces were co-visualized as multi-panel three-dimensional plots, with original experimental measurements overlaid as scatter points to explicitly denote the true data distribution.

#### Implementation and benchmarking of existing two-drug response prediction models

For benchmarking of XACT-Dual, the framework was evaluated against two state-of-the-art two-drug response prediction models: DD-PRiSM^37^ and ComboLTR^36^. Both models were implemented utilizing their official source code repositories. To ensure a standardized comparison of architectural feature extraction capabilities in the combinatorial setting, the input data modalities were harmonized across the evaluated models. Specifically, because the original ComboLTR architecture is designed to utilize multi-omics data, its input processing pipeline was modified to restrict the model to transcriptomic profiles and chemical structural information, ensuring direct comparability with XACT-Dual. Benchmarking was conducted under 10-fold cross-validation applied to both random split and leave-cells-out conditions, consistent with the monotherapy evaluation framework.

### Model interpretability analysis

#### Transcriptomic feature attribution via SHAP

SHAP (SHapley Additive exPlanations) values were computed using the DeepExplainer approximation algorithm^25^, which is designed for efficient attribution in deep neural networks. A custom model wrapper was implemented that fixed the drug feature input to the target compound while treating the normalized enrichment score vector of each cell line as the sole variable input, isolating the contribution of pathway-level transcriptomic features independent of drug structural variability. A background reference distribution was established by randomly sampling cell lines from the training set, and SHAP values were computed for a held-out subset of 50 test-set cell lines. Global feature importance was summarized by ranking pathway signatures according to their mean absolute SHAP value across the evaluated cell lines.

#### Structural mechanism analysis with GNNExplainer

GNNExplainer was applied as a perturbation-based local interpretability method for the graph neural network component to elucidate the molecular substructures driving sensitivity predictions at the atomic level. A sensitivity-filtered aggregation strategy was employed: for each target compound, all cell lines in the test set were ranked by their predicted IC_50_ values, and analysis was restricted to the 15 cell lines exhibiting the lowest predicted IC_50_. For each of these 15 samples, an edge mask was optimized over 200 epochs to identify the atomic bonds most critical to model predictions, and the resulting masks were averaged across samples to produce a consensus importance score for each atomic bond. Consensus importance scores were visualized using RDKit with atomic bonds color-coded by mean importance score.

### XACT-Clinical development and validation using TCGA data

#### Cohort selection

RNA-seq gene expression data and clinical treatment records were obtained from the open-access TCGA Pan-Cancer dataset via cBioPortal^61^. The cohort was restricted to patients who had received combination chemotherapy consisting of two or more antineoplastic agents and for whom both tumor transcriptomic profiles and clinical outcome annotations were available, yielding a final analysis cohort of 963 patients encompassing 2,779 drug pair records. For patients administered three or more drugs, all possible pairwise drug combinations were enumerated and each pair was assigned the clinical outcome label of the overall regimen.

#### Domain adaptation

To adapt XACT-Dual to the clinical prediction task, the final regression layer was replaced with a binary classification head and optimized on TCGA treatment outcome data, yielding XACT-Clinical. Clinical responses were defined according to the RECIST classification framework^39^, with patients achieving Complete Response (CR) or Partial Response (PR) designated as responders (label 0) and those exhibiting Stable Disease (SD) or Progressive Disease (PD) designated as non-responders (label 1). The drug concentration input was fixed to the pharmacokinetically relevant maximum plasma concentration (C_max_) of each drug (Supplementary Table 7). Model performance was evaluated through 10-fold cross-validation partitioned at the patient level, preventing data leakage arising from the within-patient correlation structure of the augmented dataset.

#### *In silico* virtual screening

A computational virtual screening pipeline was developed to systematically identify optimal combination partners for standard-of-care anchor drugs across patient cohorts. For a given patient cohort, the transcriptomic profile of each patient was paired with every candidate compound in the TCGA drug library under an anchor-partner strategy, in which a standard-of-care agent was fixed as the anchor drug and all remaining compounds in the library were evaluated as candidate partners. XACT-Clinical generated a predicted resistance score for every patient-anchor-partner triplet, assembling a *N*_*pbtients*_ × *M*_*drugs*_ sensitivity matrix *S* in which each entry *S*_*i,j*_ represents the predicted resistance of patient *i* to the combination of the anchor drug with partner drug i. Candidate partners were ranked by their mean predicted resistance score across the non-responding patient subgroup, with lower scores indicating greater predicted sensitivity and higher repurposing priority.

#### Pan-cancer pharmacogenomic resistance landscape

To visualize the pan-cancer landscape of combinatorial pharmacogenomic vulnerabilities, a predicted resistance matrix was constructed encompassing 963 TCGA patients across 2,775 unique drug pairs. Patient-level predicted resistance scores for each drug combination were arranged into a two-dimensional dense matrix, with patients stratified by cancer types derived from TCGA clinical annotations. To identify patient subgroups with similar pharmacological sensitivity profiles, agglomerative hierarchical clustering was performed within each cancer type cohort comprising more than two patients.

To highlight drug pairs exhibiting cancer-type-specific vulnerabilities, combinatorial efficacy was evaluated across cancer types by computing the mean predicted resistance for each drug pair within each cancer type. Drug pairs were retained for visualization if they demonstrated a minimum cross-cancer range of 0.3 and a minimum standard deviation of 0.1 in mean resistance scores across cancer types, ensuring that only combinations exhibiting biologically meaningful differential activity were displayed. The retained drug pairs were then subjected to unsupervised hierarchical clustering across the column axis based on their pan-patient predicted resistance profiles, grouping combinations with similar therapeutic activity spectra.

All hierarchical clustering was executed using Ward’s minimum variance algorithm with Euclidean distance as the dissimilarity metric. Missing values were imputed using the global median resistance score prior to clustering to ensure matrix completeness. The completely sorted and clustered resistance matrix was visualized as a high-resolution heatmap with patients arranged along the vertical axis sorted by cancer type and drug pairs along the horizontal axis.

#### Multi-instance learning framework for multi-drug combination prediction

In the TCGA cohort, patients frequently receive regimens comprising three or more agents, each of which may be combined with other drugs in the regimen in distinct pairwise configurations. To model the efficacy of such complex multi-drug regimens holistically, the prediction task was formulated within a Multi-Instance Learning (MIL) framework^45^. Each patient is represented as a "bag", and the set of pairwise drug combinations constituting their regimen serves as the "instances" within that bag, with bag-level labels defined as 0 for responders and 1 for non-responders according to RECIST criteria.

XACT-Clinical generates a forward pass for all constituent drug pair instances within a patient bag, producing raw logits that are passed through a sigmoid activation function to yield instance-level predicted resistance scores. These instance-level scores are aggregated into a single patient-level prediction through min-pooling, such that for a patient bag containing *N* drug combination instances producing predicted scores v_i_, v_2_, …, v_n_, the final prediction is:

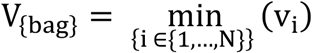

The clinical rationale for this min-pooling reflects the pharmacological principle that the overall therapeutic outcome of a multi-drug regimen is determined by its most effective constituent combination. Trainable parameters were optimized using the AdamW optimizer with a learning rate of 5×10^-5^ and a weight decay of 1×10^-4^, minimizing the Binary Cross-Entropy Loss between the min-pooled bag prediction and the ground-truth patient label. Training was conducted for 15 epochs with model selection based on ROC-AUC evaluated on a held-out validation set at the end of each epoch.

### Experimental Validation of XACT-Single predictions via cell viability assay

#### *In silico* candidate selection

To identify potential therapeutic candidates for the SaOS-2 osteosarcoma cell line, computational screening was performed using the leave-cells-out pre-trained XACT-Single model, which was applied to predict cell viability for all compounds in the GDSC library against SaOS-2, a cell line entirely absent from the training dataset. Compounds predicted to exhibit high cytotoxic efficacy were prioritized, and two mechanistically distinct agents (MG-132 and elesclomol) were selected for subsequent *in vitro* experimental validation.

#### Cell culture

The SaOS-2 human osteosarcoma cell line was maintained in complete DMEM (C-DMEM; Nacalai Tesque) in a humidified incubator at 37℃ with 5% CO_2_. For experimental preparation, cells were washed with phosphate-buffered saline (PBS), detached using Trypsin/EDTA for 5 min at 37℃, and neutralized with C-DMEM. The cell suspension was centrifuged at 200×g for 3 min, and viable cells were counted using an automated cell counter with trypan blue exclusion prior to seeding.

#### Cell viability assay

SaOS-2 cells were seeded in 96-well flat-bottom transparent microplates (Costar) in 100 μL of C-DMEM at a density of 2.5 × 10^3^ cells per well for MG-132 experiments and 1 × 10⁴ cells per well for elesclomol experiments. After an initial attachment period, MG-132 and elesclomol were added to the wells from 100 mM DMSO stock solutions (Sigma) and serially diluted to final concentrations of 0.01, 0.1, 1, and 10 μM. Negative control wells received an equivalent volume of vehicle at a final DMSO concentration of 0.1%. Cells were treated for 72h, after which 110 μL of MTT working solution diluted in C-DMEM was added to each well and incubated for 4h at 37 ° C. Formazan precipitates were dissolved by adding 100 μ L of solubilization solution and thorough mixing by pipetting. Absorbance was measured at a primary wavelength of 570 nm with a reference wavelength of 650 nm using a microplate reader (TECAN Spark). Relative cell viability was calculated as:

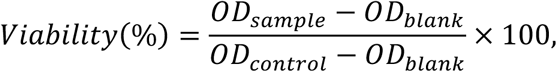

where *OD*_*sbmple*_ is the absorbance of the drug-treated wells, *OD*_*control*_ is the absorbance of the vehicle control wells, and *OD*_*blbnk*_ is the background absorbance of the medium-only wells.

#### Statistical analysis

Predictive performance in the preclinical benchmarking tasks was quantified utilizing four primary metrics: Root Mean Square Error (RMSE), Pearson Correlation Coefficient (PCC), Spearman’s Rank Correlation Coefficient (SCC), and the Coefficient of Determination (R^2^), computed utilizing the scikit-learn (version 0.24.2) and scipy (version 1.7.1) libraries in Python (version 3.9.7). Statistical significance of performance differences between XACT and the baseline models was assessed by two-sided Student’s *t*-tests implemented in the scipy.stats module.

Differences in XACT resistance scores between RECIST-defined responder and non-responder groups were assessed using the Mann-Whitney U test^62^. For patients contributing multiple drug pair scores, a single patient-level predictive score was derived by averaging across all available pairs. The cohort was then stratified into predicted-resistant and predicted-sensitive groups at the median resistance score, and overall survival was estimated using the Kaplan-Meier method^63^ with between-group differences assessed by the log-rank test. To evaluate the independence of the XACT resistance score as a prognostic factor, multivariate Cox proportional hazards regression^40^ was performed incorporating patient age, gender, clinical stage, and cancer type as covariates. Cancer types represented by fewer than 15 patients and cases with incomplete clinical annotation were excluded to ensure model stability. Statistical significance was defined as a two-sided *p*-value below 0.05 for all analyses.

## Data availability

All training data regarding monotherapy IC_50_ values, cell viability values and two-drug cell viability values, as well as clinical data used were obtained from public databases^8,35,38^.

## Acknowledgements

This work was supported by KAKENHI grants from the Japan Society for the Promotion of Science (JSPS) to H.S (23K28184, 23K18502, 24H01755, and 25H01571), JST FOREST Program to H.S. (JPMJFR242Q), as well as the Canon Foundation. We thank H. Hishinuma, T. Sakuma, and Y. Otani for critical reading of the manuscript, K. Tanaka for help with preparation of the manuscript and all laboratory members for discussion.

## Author contributions

H.S. conceived of, designed, and supervised the study. K.O. developed XACT, performed the computational study with inputs from T.I. K.O. and H.S. wrote the manuscript. All authors have read and approved the final manuscript.

