## Supplementary Tables S1-S7, Supplementary Figures S1-S4 for "Dose-dependent modeling of combinatorial drug responses stratifies patient survival and reveals therapeutic vulnerabilities in precision oncology"

**Supplementary Table 1 | Performance of XACT-Single and existing $IC_{50}$ prediction models under random split cross-validation**

| Model | RMSE | PCC | SCC | $R^2$ |
|---|---|---|---|---|
| **XACT-Single** | **1.091 ± 0.005** | **0.924 ± 0.001** | **0.912 ± 0.001** | **0.852 ± 0.001** |
| HiDRA[13] | 1.215 ± 0.007 | 0.904 ± 0.001 | 0.891 ± 0.002 | 0.817 ± 0.003 |
| PathDNN[11] | 1.279 ± 0.006 | 0.894 ± 0.001 | 0.880 ± 0.002 | 0.797 ± 0.003 |
| MLP | 1.332 ± 0.014 | 0.884 ± 0.002 | 0.869 ± 0.001 | 0.780 ± 0.003 |
| CDS[10] | 1.532 ± 0.027 | 0.843 ± 0.007 | 0.814 ± 0.011 | 0.708 ± 0.013 |

**Supplementary Table 2 | Performance of XACT-Single and existing $IC_{50}$ prediction models under leave-cells-out cross-validation**

| Model | RMSE | PCC | SCC | $R^2$ |
|---|---|---|---|---|
| **XACT-Single** | **1.267 ± 0.018** | **0.895 ± 0.002** | **0.879 ± 0.003** | **0.801 ± 0.006** |
| HiDRA[13] | 1.374 ± 0.017 | 0.876 ± 0.003 | 0.857 ± 0.004 | 0.766 ± 0.005 |
| PathDNN[11] | 1.498 ± 0.031 | 0.853 ± 0.006 | 0.835 ± 0.008 | 0.721 ± 0.012 |
| MLP | 1.491 ± 0.014 | 0.859 ± 0.003 | 0.837 ± 0.004 | 0.724 ± 0.006 |
| CDS[10] | 1.671 ± 0.016 | 0.810 ± 0.005 | 0.773 ± 0.006 | 0.653 ± 0.007 |

**Supplementary Table 3 | Performance of XACT-Single in dose-dependent cell viability prediction under random split and leave-cells-out cross-validation**

| Data split | RMSE | PCC | SCC | R2 |
|---|---|---|---|---|
| Random split | 0.1012 ± 0.0014 | 0.8871 ± 0.0028 | 0.6847 ± 0.0034 | 0.7784 ± 0.0047 |
| Leave-cells-out | 0.1372 ± 0.0032 | 0.7701 ± 0.0080 | 0.5767 ± 0.0124 | 0.5880 ± 0.0131 |

**Supplementary Table 4 | Performance of XACT-Dual and existing two-drug response prediction models under random split cross-validation**

| Model | RMSE | PCC | R2 |
|---|---|---|---|
| **XACT-Dual** | **0.072 ± 0.000** | **0.965 ± 0.001** | **0.930 ± 0.001** |
| DD-PRiSM[37] | 0.124 ± 0.002 | 0.892 ± 0.003 | 0.796 ± 0.006 |
| ComboLTR[36] | 0.211 ± 0.005 | 0.639 ± 0.022 | 0.407 ± 0.029 |

**Supplementary Table 5 | Performance of XACT-Dual and existing two-drug response prediction models under leave-cells-out cross-validation**

| Model | RMSE | PCC | R2 |
|---|---|---|---|
| **XACT-Dual** | **0.166 ± 0.014** | **0.803 ± 0.019** | **0.653 ± 0.048** |
| DD-PRiSM[37] | 0.177 ± 0.011 | 0.770 ± 0.024 | 0.575 ± 0.053 |
| ComboLTR[36] | 0.239 ± 0.017 | 0.530 ± 0.028 | 0.226 ± 0.087 |

**Supplementary Table 6 | Multivariate Cox proportional hazards regression analysis of the XACT resistance score and clinical covariates**

| **Covariate** | HR | 95% CI (lower) | 95% CI (upper) | *p*-value | *n* |
|---|---|---|---|---|---|
| XACT resistance score | 2.47 | 1.14 | 5.35 | 0.022 | 727 |
| Age | 1.02 | 1.01 | 1.03 | 0.002 | 727 |
| Gender (Female, compared to male) | 0.97 | 0.75 | 1.25 | 0.808 | 357 |
| Stage II vs. I | 0.96 | 0.71 | 1.30 | 0.795 | 294 |
| Stage III vs. I | 1.19 | 0.88 | 1.63 | 0.258 | 249 |
| Stage IV vs. I | 1.74 | 1.19 | 2.56 | 0.005 | 89 |
| Cancer type: BRCA | 0.53 | 0.36 | 0.78 | 0.001 | 174 |
| Cancer type: COADREAD | 0.92 | 0.61 | 1.40 | 0.694 | 82 |
| Cancer type: ESCA | 1.38 | 0.59 | 3.23 | 0.463 | 23 |
| Cancer type: HNSC | 2.34 | 1.13 | 4.86 | 0.023 | 15 |
| Cancer type: LUAD | 1.31 | 0.88 | 1.94 | 0.184 | 88 |
| Cancer type: LUSC | 1.12 | 0.70 | 1.79 | 0.643 | 56 |
| Cancer type: MESO | 2.76 | 1.69 | 4.50 | < 0.001 | 25 |
| Cancer type: PAAD | 2.04 | 1.21 | 3.44 | 0.007 | 35 |
| Cancer type: SKCM | 1.41 | 0.77 | 2.56 | 0.264 | 22 |

| | | | | | |
|---|---|---|---|---|---|
| Cancer type: STAD | 1.78 | 1.16 | 2.72 | 0.008 | 74 |
| Cancer type: TGCT | 0.33 | 0.17 | 0.64 | 0.001 | 52 |

HR, hazard ratio; CI, confidence interval; n, number of patients included in each comparison. Reference category for tumor stage is Stage I; reference category for cancer type is COADREAD (the most prevalent cancer type in the cohort after exclusion criteria were applied). All p-values are two-sided. Cancer type abbreviations follow TCGA nomenclature: BRCA, breast invasive carcinoma; HNSC, head and neck squamous cell carcinoma; LUAD, lung adenocarcinoma; LUSC, lung squamous cell carcinoma; MESO, mesothelioma; PAAD, pancreatic adenocarcinoma; SKCM, skin cutaneous melanoma; STAD, stomach adenocarcinoma; TGCT, testicular germ cell tumors; COADREAD, colorectal adenocarcinoma.

**Supplementary Table 7 | Maximum plasma concentration ($C_{max}$) values of anticancer drugs used in XACT-Clinical analysis**

| Drug Class | Drug Name | $C_{max}$ (μM) |
|---|---|---|
| Platinum compounds | Carboplatin | 150 |
| | Cisplatin | 10 |
| | Oxaliplatin | 2 |

| | | |
|---|---|---|
| Taxanes | Cabazitaxel | 0.27 |
| | Docetaxel | 4.6 |
| | Ixabepilone | 0.5 |
| | Paclitaxel | 15 |
| Vinca alkaloids | Vinblastine | 0.01 |
| | Vincristine | 0.01 |
| | Vinorelbine | 0.15 |
| Antimetabolites | Capecitabine | 10 |
| | Doxifluridine | 20 |
| | Fluorouracil | 50 |
| | Gemcitabine | 100 |
| | Hydroxyurea | 300 |
| | Leucovorin | 150 |
| | Methotrexate | 10 |
| | Pemetrexed | 300 |
| Anthracyclines | Doxorubicin | 2.2 |
| | Epirubicin | 10 |
| | Mitoxantrone | 1.5 |
| Alkylating agents | Carmustine | 6 |
| | Cyclophosphamide | 15 |
| | Dacarbazine | 4 |

| | | |
|---|---|---|
| | Evofosfamide | 20 |
| | Fotemustine | 20 |
| | Ifosfamide | 100 |
| | Lomustine | 1.3 |
| | Melphalan | 5 |
| | Nimustine | 5 |
| | Treosulfan | 300 |
| Topoisomerase inhibitors | Cositecan | 1 |
| | Dactinomycin | 0.001 |
| | Etoposide | 30 |
| | Irinotecan | 3 |
| | Teniposide | 75 |
| | Topotecan | 0.15 |
| HDAC inhibitors | Belinostat | 53 |
| | Vorinostat | 4.5 |
| PARP inhibitors | Veliparib | 11 |
| Kinase inhibitors | Axitinib | 0.07 |
| | Dabrafenib | 4.3 |

| | | |
|---|---|---|
| | Erlotinib | 4.1 |
| | Everolimus | 0.023 |
| | Imatinib | 6.9 |
| | Lapatinib | 4.2 |
| | Mibefradil | 0.3 |
| | Palbociclib | 0.22 |
| | Pazopanib | 130 |
| | Regorafenib | 6.6 |
| | Ridaforolimus | 0.028 |
| | Sorafenib | 14 |
| | Sunitinib | 3 |
| | Tivozanib | 0.044 |
| | Trametinib | 0.036 |
| | Vandetanib | 1.4 |
| | Vemurafenib | 116 |
| Microtubule inhibitors | Eribulin | 0.002 |
| | Trabectedin | 0.01 |
| Other cytotoxics | Bleomycin | 1 |
| | Mitomycin | 2 |
| | Streptozocin | 2 |

| | | |
|---|---|---|
| Hormone modulators | Dexamethasone | 2.5 |
| | Isotretinoin | 5 |
| | Tamoxifen | 1 |
| Other / limited PK data | Didox | 10 |
| | Mitotane | 40 |
| | Polysaccharide-k | 10 |
| | Procarbazine | 12 |
| | Temozolomide | 40 |
| | Zoledronic acid | 5 |

$C_{max}$ values represent the maximum plasma concentration of each drug at standard clinical dosing, used as the pharmacokinetically relevant concentration input for XACT-Clinical predictions. Values were compiled from published pharmacokinetic studies, prescribing information, and clinical trial reports. Where multiple dosing regimens or patient populations were reported, the value corresponding to standard first-line dosing in adult patients was selected. For drugs with limited pharmacokinetic data (categorized as "Other / limited PK data"), values represent best available estimates from the literature. Drug classes follow standard oncological pharmacological classifications. $C_{max}$ values are reported in micromolar (μM) units.

### Supplementary Figure Legends

**Supplementary Fig. 1 | Transcriptomic and structural interpretability analyses for the MEK inhibitor trametinib**

**(A)** Transcriptomic determinants of trametinib response identified by SHAP analysis. SHAP summary plot displaying the ten pathway-level features exerting the greatest influence on the predicted $IC_{50}$ for trametinib. Each point represents an individual cell line, with color encoding the normalized enrichment score of the corresponding pathway signature (red: high activity; blue: low activity). The horizontal axis quantifies the direction and magnitude of each feature's contribution to the model output, with negative values indicating features associated with predicted sensitivity (lower $IC_{50}$) and positive values indicating features associated with predicted resistance (higher $IC_{50}$). Features are ranked in descending order by mean absolute SHAP value. The autonomous prioritization of KRAS and RAF signaling as one of the dominant transcriptomic determinants of trametinib sensitivity is consistent with the canonical role of MEK as a critical downstream effector of the RAS–RAF–MEK–ERK cascade, confirming that XACT-Single has captured pharmacologically valid genotype-sensitivity relationships without explicit mechanistic supervision. **(B)** Structural basis of trametinib potency identified by GNNExplainer. Atomic importance map for trametinib generated by aggregating GNNExplainer edge masks across the cell lines with the lowest predicted $IC_{50}$, representing the cellular contexts in which the drug is predicted to be most efficacious. Bond importance is encoded on a continuous red color scale, with darker red indicating higher consensus importance scores. The model prioritizes the pyridopyrimidine scaffold and the fluorobenzyl-linked region as the dominant structural determinants of potency. The convergence of high transcriptomic importance on MAPK pathway signatures and high structural importance on the allosteric pharmacophore of trametinib demonstrates that XACT-Single has learned pharmacologically valid structure-activity relationships without explicit mechanistic supervision.

**Supplementary Fig. 2 | Reconstruction of dose-dependent cell viability curves across diverse cancer types and drug classes**

**(A-I)** Predicted and experimentally observed dose-response curves for nine representative drug-cell line pairs spanning a broad range of cancer types, drug classes, and pharmacodynamic response profiles: MCF7 treated with tamoxifen (Breast, ER+) **(A)**, MCF7 treated with docetaxel (Breast, ER+) **(B)**, A498 treated with sorafenib (Kidney, RCC) **(C)**, A375 treated with vorinostat (Skin, Melanoma) **(D)**, OCI-AML3 treated with 5-fluorouracil (Blood, AML) **(E)**, HCT-116 treated with irinotecan (Colorectal) **(F)**, U-2 OS treated with vorinostat (Bone, Osteosarcoma) **(G)**, HCC70 treated with buparlisib (Breast, TNBC) **(H)**, and A549 treated with cediranib (Lung, NSCLC) **(I)**. In each panel, the horizontal axis represents the drug concentration in micromolar units on a logarithmic scale and the vertical axis represents the fractional cell viability normalized to untreated controls.

Experimentally measured viability values are shown as blue circles and connected by a smoothing spline, and XACT-Single predictions are shown as red diamonds fitted with a Hill function to capture the sigmoidal pharmacodynamic relationship between dose and cellular response. ER, estrogen receptor; RCC, renal cell carcinoma; AML, acute myeloid leukemia; TNBC, triple-negative breast cancer; NSCLC, non-small cell lung cancer.

**Supplementary Fig. 3 | Reconstruction of three-dimensional dose-response surfaces for representative two-drug combinations**

Three-dimensional dose-response surfaces illustrating the combined effect of two co-administered therapeutic agents on fractional cell viability across two cancer cell line-drug pair combinations: OVCAR-8 treated with paclitaxel plus daunorubicin (Ovary, High-grade Serous Ovarian Cancer) **(A)** and U251 treated with temozolomide plus etoposide (Brain, glioblastoma multiforme) **(B)**. For each combination, the XACT-Dual predicted surface is shown in the left panel and the corresponding experimentally observed surface is shown in the right panel. Red circles indicate measured viability values at specific drug concentration pairs, with the continuous surface generated by interpolation between measured points. The x- and y-axes represent the concentrations of the two drugs on logarithmic scales in micromolar units, and the z-axis represents fractional cell viability normalized to untreated controls. Surface color encodes the viability level on a continuous scale from 0 (dark purple, complete cell death) to 1.0

(yellow, full survival), as indicated by the shared color bar below each panel pair. The close correspondence between predicted and experimentally observed surfaces across combinations exhibiting qualitatively distinct synergy profiles demonstrates that XACT-Dual accurately reconstructs the full two-dimensional dose-response manifold across diverse pharmacological mechanisms.

**Supplementary Fig. 4 | Virtual screening analyses in sarcoma and pancreatic adenocarcinoma**

**(A)** Recapitulation of clinical outcomes in pancreatic adenocarcinoma. Box plots displaying XACT resistance scores for TCGA pancreatic adenocarcinoma (TCGA-PAAD) patients stratified by RECIST-defined clinical outcome. Non-responders exhibited higher resistance scores than responders. **(B)** *In silico* gemcitabine anchor-partner screening for pancreatic adenocarcinoma. Box plots displaying XACT resistance scores for the three highest-ranked alternative combination partners for gemcitabine across non-responding PAAD patients, with the standard-of-care partner paclitaxel shown for reference. Statistical significance of pairwise comparisons against paclitaxel was assessed by two-sided $t$-test (*$p < 0.05$; ***$p < 0.001$). **(C)** *In silico* alternative first-line screening for sarcoma. Paired plots comparing the XACT resistance score of each non-responding TCGA-SARC patient's actual first-line treatment against three computationally nominated alternative regimens exhibiting substantially lower predicted resistance across the non-responding subgroup ($p = 0.0002$). Lines connect individual patients across conditions, illustrating the consistent reduction in predicted resistance achieved by the nominated regimens relative to the prescribed treatment. **(D)** *In silico* alternative first-line screening for pancreatic adenocarcinoma. Paired plots comparing the XACT resistance score of each non-responding TCGA-PAAD patient's actual first-line treatment against three computationally nominated alternative regimens. Lines connect individual patients across conditions.

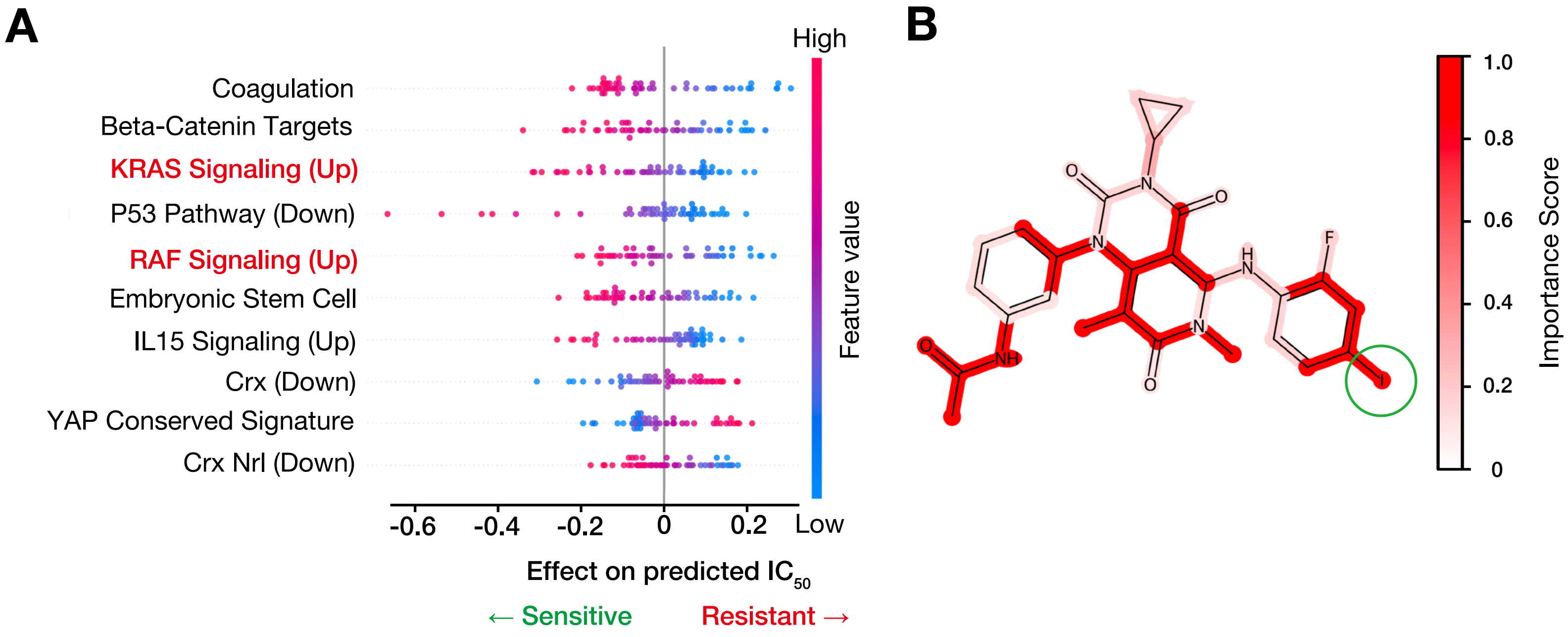


**Ota et al., Supplementary Figure 1**

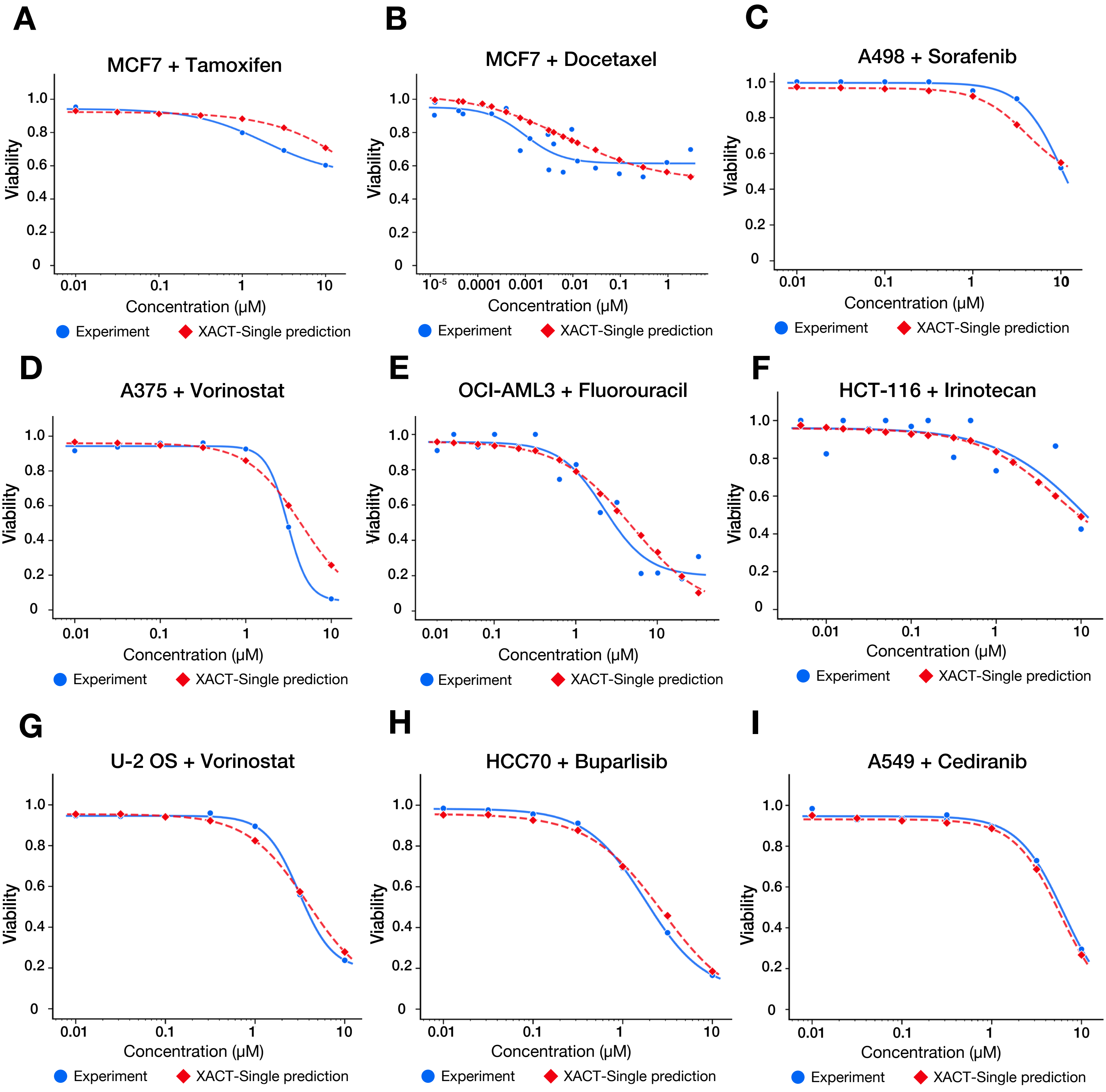


**Ota et al., Supplementary Figure 2**

**A**

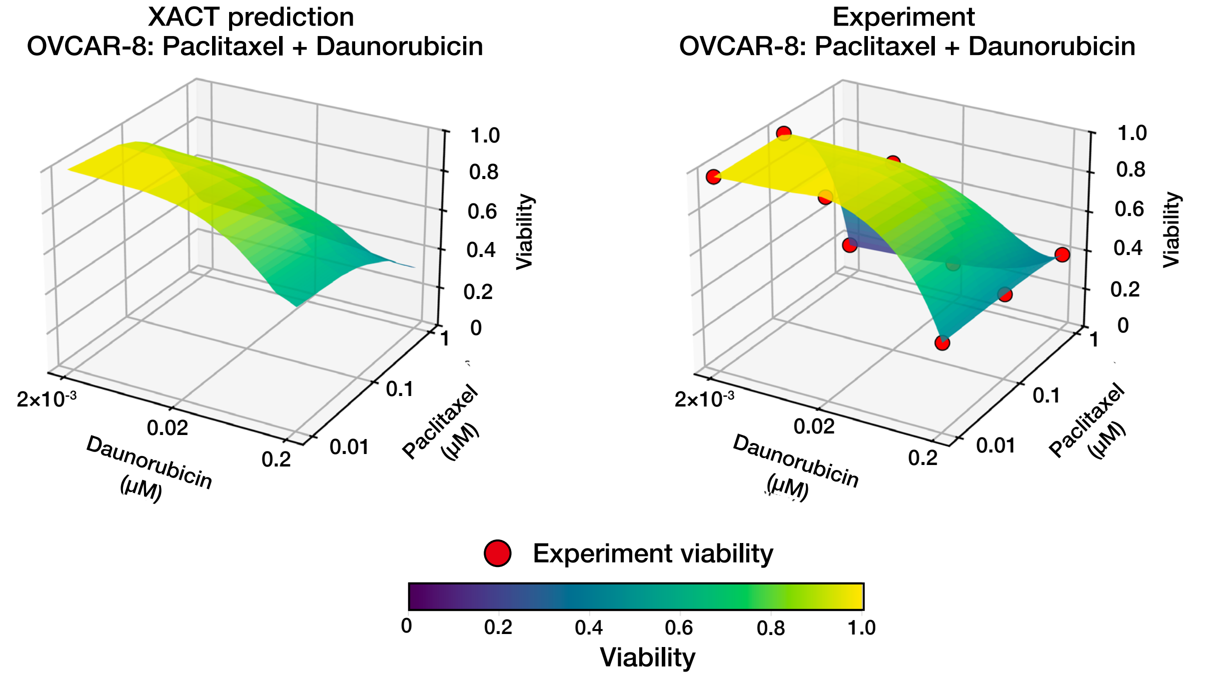


**B**

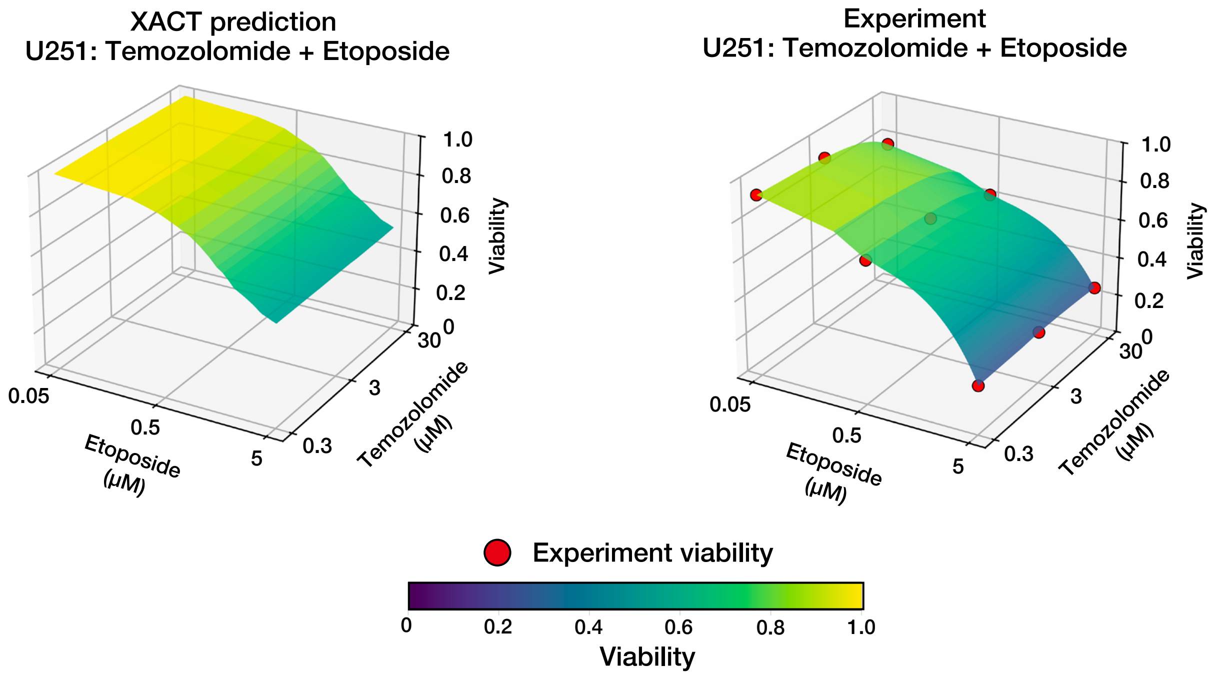


**Ota et al., Supplementary Figure 3**

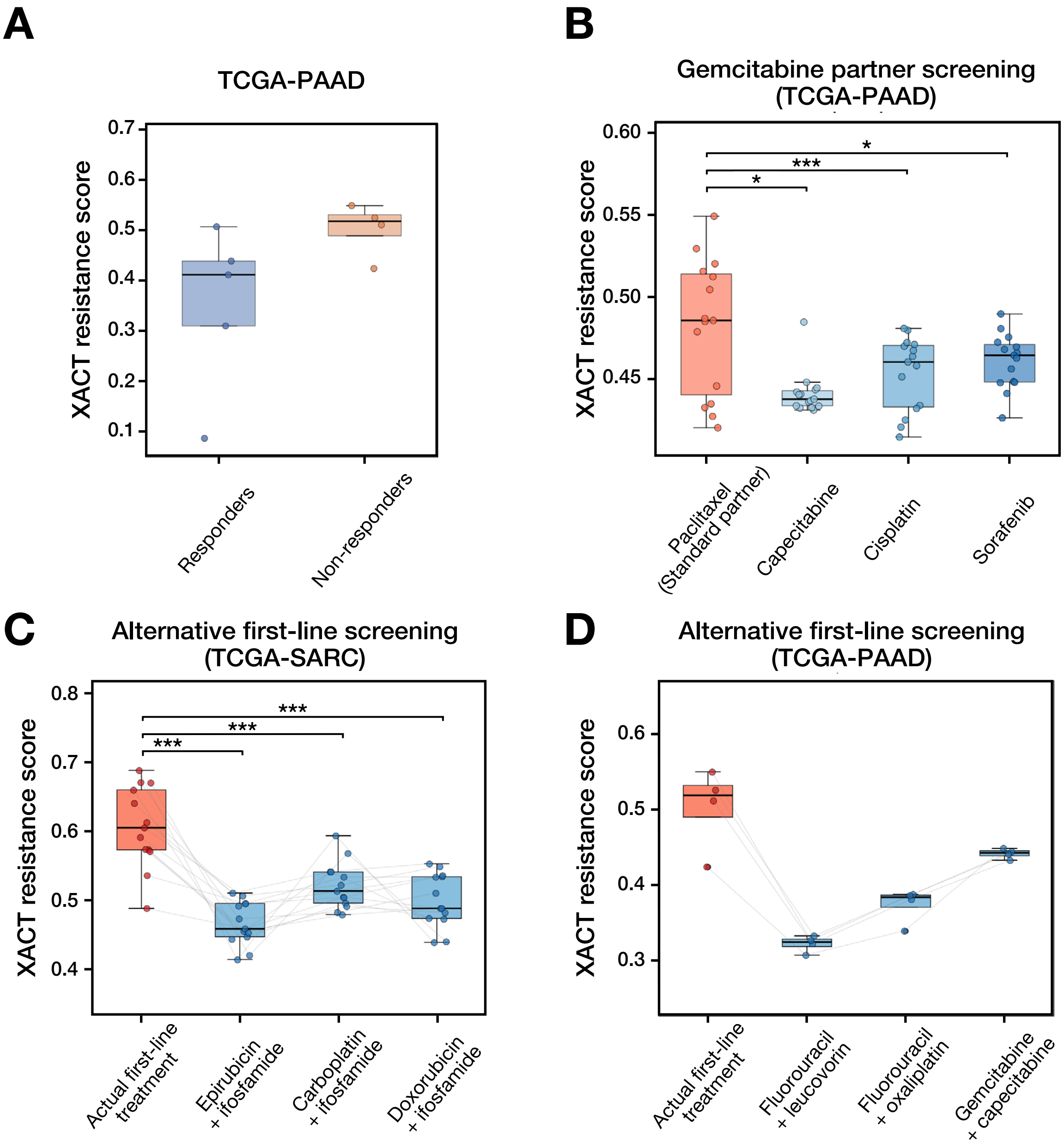


**Ota et al., Supplementary Figure 4**
